# Defining the RBPome of T helper cells to study higher order post-transcriptional gene regulation

**DOI:** 10.1101/2020.08.20.259234

**Authors:** Kai P. Hoefig, Alexander Reim, Christian Gallus, Elaine H. Wong, Gesine Behrens, Christine Conrad, Meng Xu, Taku Ito-Kureha, Kyra Defourny, Arie Geerlof, Josef Mautner, Stefanie M. Hauck, Dirk Baumjohann, Regina Feederle, Matthias Mann, Michael Wierer, Elke Glasmacher, Vigo Heissmeyer

**Affiliations:** Research Unit Molecular Immune Regulation, Helmholtz Center Munich, Munich, Germany; Department of Proteomics and Signal Transduction, Max-Planck-Institute of Biochemistry, Munich, Germany; Institute of Diabetes and Obesity, Helmholtz Center Munich, Munich, Germany; Institute for Immunology at the Biomedical Center, Ludwig-Maximilians-Universität München, Planegg-Martinsried, Germany; Institute of Structural Biology, Helmholtz Center Munich, Neuherberg, Germany; Research Unit Gene Vectors, Helmholtz Center Munich & Children’s Hospital, TU Munich, Germany; Research Unit Protein Science, Helmholtz Center Munich, Munich, Germany; Monoclonal Antibody Core Facility and Research Group, Institute for Diabetes and Obesity, Helmholtz Center Munich, Neuherberg, Germany; Roche Pharma Research and Early Development, Large Molecule Research, Roche Innovation Center Munich, Penzberg, Germany; Department of Biomolecular Health Sciences, Utrecht University, Utrecht, the Netherlands; Medical Clinic III for Oncology, Immuno-Oncology and Rheumatology University Hospital Bonn, University of Bonn, Bonn, Germany; Proteomics Research Infrastructure, University of Copenhagen, Copenhagen, Denmark

## Abstract

Post-transcriptional gene regulation is complex, dynamic and ensures proper T cell function. The targeted transcripts can simultaneously respond to various factors as evident for *Icos*, an mRNA regulated by several RNA binding proteins (RBPs), including Roquin. However, fundamental information about the entire RBPome involved in post-transcriptional gene regulation in T cells is lacking. Here, we applied global RNA interactome capture (RNA-IC) and orthogonal organic phase separation (OOPS) to human and mouse primary T cells and identified the core T cell RBPome. This defined 798 mouse and 801 human proteins as RBPs, unexpectedly containing signaling proteins like Stat1, Stat4 and Vav1. Based on the vicinity to Roquin-1 in proximity labeling experiments, we selected ∼50 RBPs for testing coregulation of Roquin targets. Induced expression of these candidate RBPs in wildtype and Roquin-deficient T cells unraveled several Roquin-independent contributions, but also revealed Celf1 as a new Roquin-1-dependent and target-specific coregulator of *Icos*.

**One sentence statement:** We provide an atlas of RNA-binding proteins in human and mouse T helper cells as a resource for studying higher order post-transcriptional gene regulation.

---

T lymphocytes as central entities of the adaptive immune system must be able to make critical cell fate decisions fast. To exit quiescence, commit to proliferation and differentiation, exert effector functions or form memory they strongly depend on programs of gene regulation. Accordingly, they employ extensive post-transcriptional regulation through RBPs and miRNAs or 3’ end oligo-uridylation and m6A RNA modifications. These RBPs, or RBPs that recognize the modifications, directly affect the expression of genes by controlling mRNA stability or translation efficiency ^1, 2, 3, 4, 5^. The studies in T helper cells have focused on a small number of RNA-binding proteins, including HuR and TTP/Zfp36l1/Zfp36l2 ^6, 7, 8, 9^, Roquin-1/2 ^10, 11^ and Regnase-1/4 ^12, 13^ as well as some miRNAs like miR-17∼92, miR-155, miR-181, miR-125 or miR-146a ^14^. Moreover, first evidence for m6A RNA methylation in this cell type has been provided ^4^. Underscoring the relevance for the immune system, loss-of-function of these factors has often been associated with profound alterations in T cell development or functions which caused immune-related diseases ^14, 15, 16, 17^. Intriguingly, many key factors of the immune system have acquired long 3’-UTRs enabling their regulation by multiple, and often overlapping sets of post-transcriptional regulators ^18^. Some RBPs recruit additional co-factors as for example Roquin binding with Nufip2 to RNA ^19^ and some of them have antagonistic RBPs like HuR and TTP ^20^ or Regnase-1 and Arid5a ^21^. Such functional or physical interactions together with interdependent binding to the transcriptome create enormous regulatory potential. The challenge is therefore to integrate our current knowledge about individual RBPs into concepts of higher order gene regulation that reflect the interplay of different, and ideally of all cellular RBPs.

A prerequisite for studying higher order post-transcriptional networks is to know cell type specific RBPomes to account for differential expression and RBP plasticity. To this end several global methods have been developed over the last decade, revealing a growing number of RBPs that may even exceed recent estimates of ∼7.5% of the human proteome ^22^. RNA interactome capture (RNA-IC) ^23^ is one widely used, unbiased technique, however, it is constricted by design, exclusively identifying proteins binding to polyadenylated RNAs. In contrast, orthogonal organic phase separation (OOPS) analyzes all UV-crosslinked protein-RNA adducts from interphases after organic phase separation.

The interactions of RBPs with RNA typically involve charge-, sequence- or structure-dependent interactions and to date over 600 structurally different RNA-binding domains (RBD) have been identified in canonical RBPs of the human proteome ^22^. However, global methods also identified hundreds of non-canonical RBPs, which oftentimes contained intrinsically disordered regions (IDRs). Surprisingly, as many as 71 human proteins with well-defined metabolic functions were found to interact with RNA ^24^ introducing the concept of “moonlighting”. Depending on availability from their “day job” in metabolism such proteins also bear the potential to impact on RNA regulation. Recent large-scale approaches have increased the number of EuRBPDB-listed human RBPs to currently 2949 ^25^, suggesting that numerous RNA/RBPs interactions and cell-type specific gene regulations have gone unnoticed so far.

As a first step towards a global understanding of post-transcriptional gene regulation we experimentally defined all proteins that can be crosslinked to RNA in T helper cells. RNA-IC and OOPS identified 310 or 1200 proteins in primary CD4^+^ T cells interacting with polyadenylated transcripts or all RNA species, respectively. Importantly, this dataset now enables the study of higher order gene regulation by determining for example how the cellular RBPs participate or intervene with post-transcriptional control of targets by individual RBPs.

## Results

### Icos exhibits simultaneous and temporal regulation through several RBPs

A prominent example for complex post-transcriptional gene regulation is the mRNA encoding inducible T-cell costimulator (Icos). It harbors a long 3*’*-UTR, which responds in a redundant manner to Roquin-1 and Roquin-2 proteins, is repressed by Regnase-1 and microRNAs, but also contains TTP binding sites ^7, 10, 11, 13, 26, 27, 28^. Moreover, the Icos 3*’*-UTR was proposed to be modified by m6A methylation ^29^, which could either attract m6A-specific RBPs with YTH domains ^30^, recruit or repel other RBPs ^31^, or interfere with base-pairing and secondary structure- or miRNA/mRNA-duplex formation ^32^. Because of transcriptional and post-transcriptional regulation, Icos expression exhibits a hundred-fold upregulation on the protein level during T cell activation, which then declines after removal of the TCR stimulus (**Fig. 1**). To determine the temporal impact on Icos expression by Roquin, Regnase-1, m6A and miRNA regulation we analyzed inducible, CD4-specific inactivation of Roquin-1 together with Roquin-2 (*Rc3h1-2*), Regnase-1 (*Zc3h12a*) or Wtap, an essential component of the m6A methyltransferase complex ^33^, as well as Dgcr8, which is required for pre-miRNA biogenesis ^34^. To this end, we performed tamoxifen gavage on mice expressing a Cre-ERT2 knockin allele in the CD4 locus ^35^ together with the floxed, Roquin paralogs encoding, *Rc3h1* and *Rc3h2* alleles (**Fig. 1a-c**), Regnase-1 encoding *Zc3h12a* alleles (**Fig. 1d-f**) as well as *Wtap*, (**Fig. 1g-i**) and *Dgcr8* alleles, (**Fig. 1j-l**). We isolated CD4^+^ T cells from these mice and expanded them for five days. Confirming target deletion on the protein level (**Fig. 1c, f, i, l**) we determined a strong negative effect on Icos expression by Roquin and Regnase-1 on days 2-5 (**Fig. 1a-b** and **d-e**), a moderate positive effect of Wtap on days 2-5 (**Fig. 1g-h**), and only a small effect of Dgcr8, with an initial tendency of negative (day 1) and later on positive effects (days 4-5) (**Fig. 1j-k**). We next asked whether T cell activation affects the expression levels of known regulators of Icos, as well as other RBPs, to install temporal compartmentalization. To do so, we monitored the expression of a panel of RBPs in mouse CD4^+^ T cells over the same time course (**Fig. 1m-q**). Indeed, we observed fast upregulation of RBPs as determined with pan-Roquin, Nufip2 ^19^, Fmrp, Fxr1, Fxr2, pan-TTP/Zfp36 7, pan-Ythdf (**Supplementary Fig. 1a**), or Celf1 specific antibodies, but also slower accumulation as determined with Regnase-1 or Rbms1 specific antibodies (**Fig. 1m, o**). There was also downregulation of RBPs as shown with pan-AGO ^36^ and Cpeb4 specific antibodies (**Fig. 1m-n and p**). Of note, we also observed signs of post-translational regulation showing incomplete or full cleavage of Roquin or Regnase-1 proteins ^10, 13^, respectively (**Fig. 1m**), and the induction of a slower migrating band for Celf1, likely phosphorylation ^37^ (**Fig. 1q**). Factors with the potential to cooperate, as involved for Roquin and Nufip2^19^ or Roquin and Regnase-1^10^, showed overlapping temporal regulation (**Fig. 1m**), suggesting reinforcing effects on the Icos target. Together these data indicate that mRNA targets respond to simultaneous inputs from several RBPs, which are aligned by dynamic expression and post-translational regulation to orchestrate redundant, cooperative and antagonistic effects into a higher order regulation.

**Figure 1:**
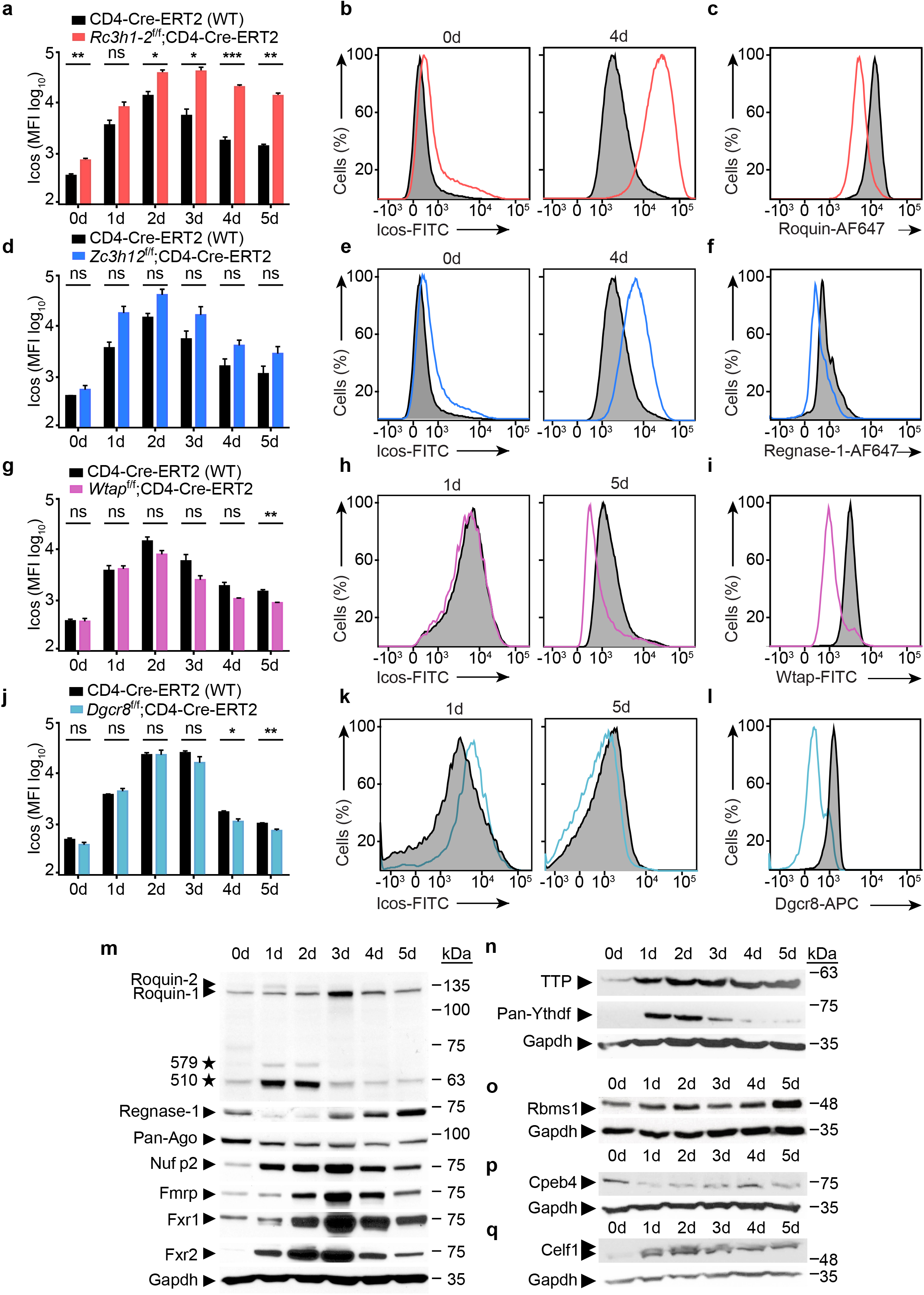
Icos responds to inputs from several post-transcriptional regulators. **a, d, g, j,** Bar diagrams of Icos mean fluorescence intensities (MFI) over a five-day period of T cell activation illustrating changes induced by the CD4^+^-specific, induced depletion of the respective genes. Significance was calculated using the unpaired t-test (two-tailed) with n=3. **b, e, h, k,** Representative histograms of Icos expression at the specified days. **c, f, i, l,** Histograms demonstrating successful depletion of the respective target proteins. **o-r,** Western blots showing patterns of dynamic RBP regulation after CD3 and CD28 CD4^+^ T cell activation. ns = not significant; * significant (p value = 0.01 – 0.05); ** very significant (p value = 0.001 – 0.01); *** very significant (p value = < 0.001).

### The polyadenylated RNA-bound proteome of mouse and human T helper cells

To decipher this post-transcriptional network we set out to determine the RNA-binding protein signature in primary mouse and human CD4^+^ T cells. To identify mRNA-binding proteins we first performed mRNA capture experiments on CD4^+^ T cells expanded under T_H_0 culture conditions (**Fig. 2a**). Pull-down with oligo-dT beads enabled the enrichment of mRNA-bound proteins as determined by silver staining, which were increased in response to preceding UV irradiation of the cells (**Supplementary Fig. 1b**). Reverse transcription quantitative PCR (RT-qPCR) confirmed the recovery of specific mRNAs, such as for the housekeeping genes *Hprt* and *β-actin*. Both mRNAs were enriched at least 2-3 fold after UV crosslink, but there was no detection of non-polyadenylated 18S rRNA (**Supplementary Fig. 1c**). Focusing on protein recovery, we determined greatly enriched polypyrimidine tract-binding protein 1 (Ptbp1) RBP compared to the negative control β-tubulin in mRNA capture experiments using the EL-4 thymoma cell line (**Supplementary Fig. 1d**). Next we performed mass spectrometry (MS) on captured proteins from murine and human T cells (**Fig. 2b-c**). Quantifying proteins bound to mRNA in crosslinked (CL) versus non-crosslinked (nCL) samples we defined a total of 312 mouse (**Fig. 2b**) and 308 human mRNA-binding proteins (mRBPs) (**Fig. 2c**) with an overlap of ∼70% (**Supplementary Table 1**), which is in concordance with the overlap of all listed RBPs for these two species in the eukaryotic RBP database (http://EuRBPDB.syshospital.org). Gene ontology (GO) analysis identified the term ‘mRNA binding’ as most significantly enriched (**Fig. 2d**). The top ten GO terms in mouse were also strongly enriched in the human dataset with comparable numbers of proteins assigned to the individual GO terms in both species (**Fig. 2d**). RBPs not only bind RNA by classical RBDs, but RNA-interactions can also map to intrinsically disordered regions (IDRs) ^38^. Furthermore, low complexity regions (LCRs) have also been reported to be overrepresented in RBPs ^23^. Indeed, IDRs (**Fig. 2e**) and LCRs (**Supplementary Fig. 1e**) were strongly enriched protein characteristics of mouse and human RNA-IC-identified RBPomes. We next wondered how much variation exists in the composition of RBPs between different T helper cell subsets. Repeating RNA-IC experiments in mouse and human iTreg cells (**Supplementary Fig. 2a**), we find an overlap of 96% or 90% with the respective mouse or human iTreg with Teff RBPomes, suggesting that the same RBPs bind to the transcriptome in different T helper cell subsets (Table 1). Nevertheless, 47 or 48 proteins were exclusively identified in mouse or human effector T cells, respectively, and 10 or 28 proteins were only found in mouse or human Treg cells, respectively (**Supplementary Fig. 2b**). Although these differences could indicate the existence of subset-specific RBPs, they could also be related to an incomplete assessment of RBPs by the RNA-IC technology.

**Figure 2:**
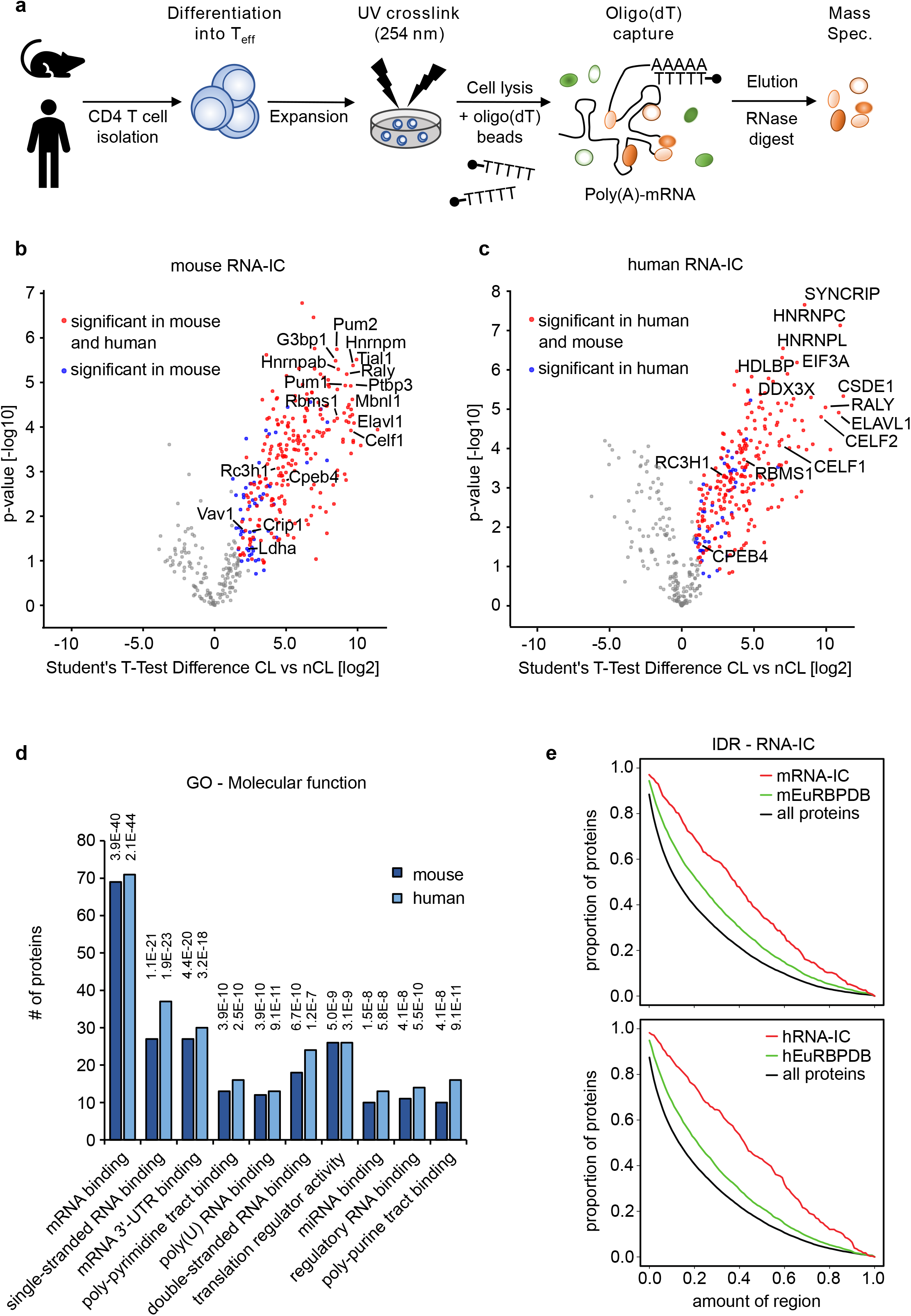
The CD4^+^ T cell RBPome of polyadenylated RNAs. **a,** Schematic illustration of the RNA-interactome capture (RNA-IC) method that was carried out to identify RBPs from mouse and human CD4^+^ T cells. **b, c,** Volcano plot showing the −log10 p-value plotted against the log2 fold-change comparing the RNA-capture from crosslinked (CL) mouse CD4 T cells (b) or human CD4^+^ T cells (c) versus the non-crosslinked (nCL) control. Red dots represent proteins significant at a 5% FDR cut-off level in both mouse and human RNA-capture experiments and blue dot proteins were significant only in mouse or human, respectively. **d,** Enrichment analysis of GO Molecular Function terms of significant proteins in mouse or human RNA capture data. The 10 most enriched terms in mouse (dark blue) and the respective terms in human (light blue) are shown. The y-axis represents the number of proteins matching the respective GO term. Numbers above each term depict the adjusted p-value after Benjamini-Hochberg multiple testing correction. **e,** Distribution of IDRs in all Uniprot reviewed protein sequences (black line), in proteins of the mouse EuRBPDB database (green line) and in proteins significant in the mouse RNA-IC experiment (red line). The same plot is shown for human data at the bottom. According to Kolmogorov-Smirnov testing the IDR distribution differences between RNA-IC (red lines) and all proteins (black lines) are highly significant in mouse and man and reach the smallest possible p-value (p<2.2×10^−16^).

### The global RNA-bound proteome of mouse and human T helper cells

We therefore employed a second RNA-centric method of orthogonal organic phase separation (OOPS) to extend our definition of the RBPome and cross-validate our results. Similar to interactome capture, OOPS preserves cellular protein/RNA interactions by UV crosslinking of intact cells. The physicochemical properties of the resulting aducts direct them towards the interphase in the organic and aqueous phase partitioning procedure (**Fig. 3a**). Following several cycles of interphase transfer and phase partitioning, RNase treatment releases RNA-bound proteins into the organic phase, making them amenable to mass spectrometry ^39, 40^. Evaluating the method, we investigated RNA and proteins from purified interphases derived from CL and nCL MEF cell samples. RNAs with crosslinked proteins purified from interphases hardly migrated into agarose gels, but regained normal migration behavior after protease digest, as judged from the typical 18S and 28S rRNA pattern (**Supplementary Fig. 3a**). Conversely, known RBPs like Roquin-1 and Gapdh appeared after crosslinking in the interphase and could be recovered after RNase treatment from the organic phase (**Supplementary Fig. 3b**). Utilizing the same cell numbers and culture conditions of T cells, this method identified in total 1255 and 1159 significantly enriched RBPs for mouse or human T cells, respectively, when comparing CL and nCL samples (**Supplementary Table 1**). The overlap between both organisms was 55% (**Fig. 3b**) and 60% (**Fig. 3c**) in relation to the individual mouse and human RBPomes. Although glycosylated proteins are known to also accumulate in the interphase ^39^, we experimentally verified that they did not migrate into the organic phase after RNase treatment (**Supplementary Fig. 3c-d**). Analyzing OOPS-derived RBPomes for gene ontology enrichment using the same approach as for RNA-IC the GO term ‘mRNA binding’ was again most significantly enriched in mouse and man (**Fig. 3d**). The top 10 GO terms were RNA related and six of them overlapped with those identified for RNA-IC-derived RBPomes. High similarity between mouse and human RBPomes becomes apparent by the similarity in all GO categories, including ‘molecular function’ (**Fig. 3d**), ‘biological process’ and ‘cellular component’ (**Supplementary Fig. 4**). Although our OOPS approach exceeded by far the quantity of RNA-IC identified RBPs, the number of ∼1200 RBPs well matched published RBPomes of HEK293 (1410 RBPs), U2OS (1267 RBPs) and MCF10A (1165 RBPs) cell lines ^39^.

**Figure 3:**
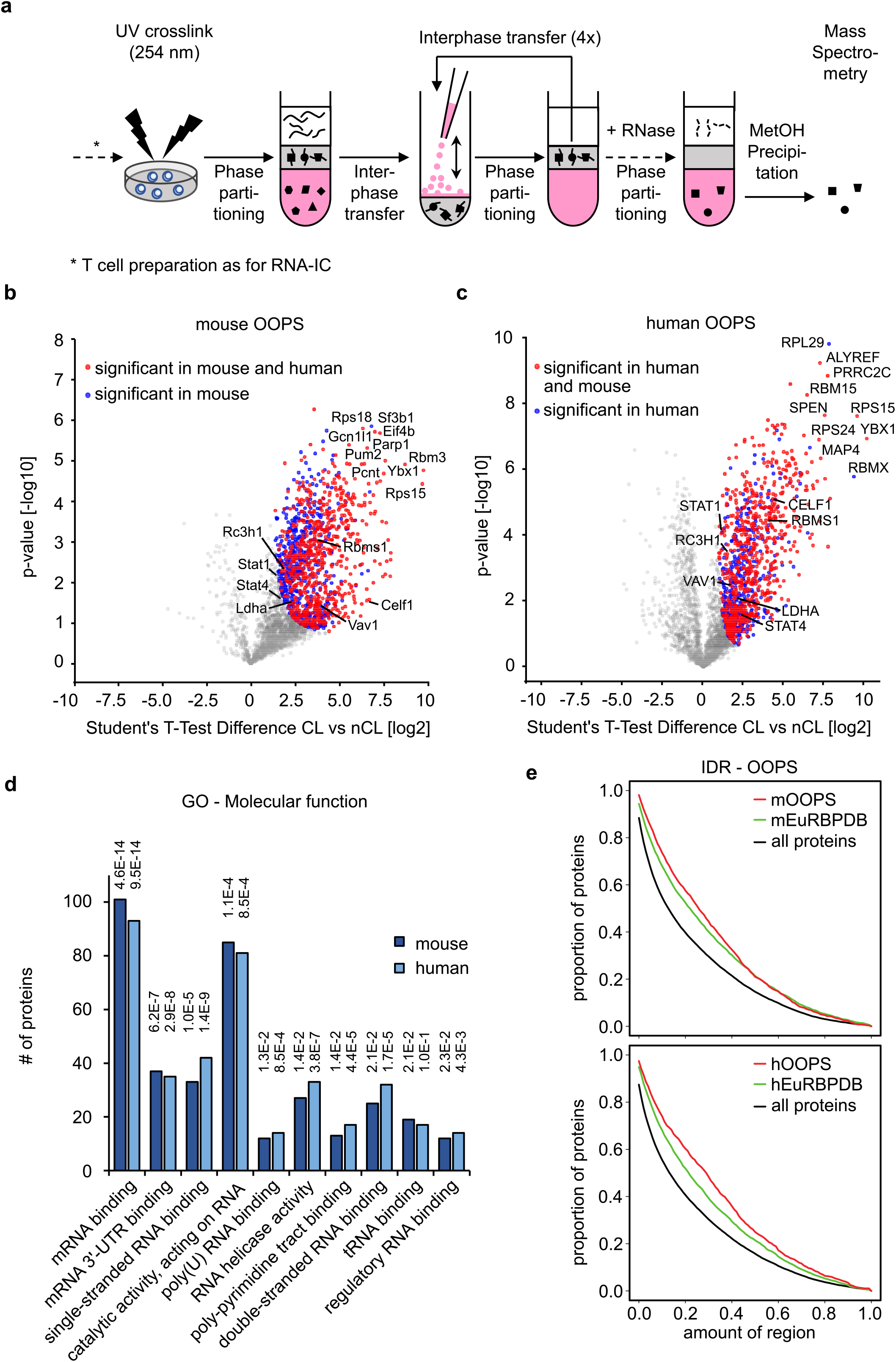
The global RNA-bound proteome of CD4^+^ T cells. **a,** Schematic overview of the OOPS method ^39^ with phase partitioning cycles increased to five. **b-c,** Volcano plots showing the −log10 p-value plotted against the log2 fold-change comparing the organic phase after RNase treatment of the interphase of OOPS experiments of crosslinked mouse CD4^+^ T cells (b) or human CD4^+^ T cells (c) versus the same non-crosslinked sample. Red dots represent proteins significant at a 5% FDR cutoff level in both mouse and human OOPS experiments and blue dots proteins significant only in mouse or human, respectively. **d,** Enrichment analysis of GO Molecular Function terms of significant proteins in mouse or human OOPS data. Enriched terms are depicted exactly as described for RNA-capture data in Figure 2d. **e,** Distribution of intrinsically disordered regions in all Uniprot reviewed protein sequences (black line), in proteins in the mouse EuRBPDB database (green) and in proteins significant in the mouse OOPS data (red line). The same plot is shown for human data at the bottom. According to Kolmogorov-Smirnov testing the IDR distribution differences between RNA-IC (red lines) and all proteins (black lines) are highly significant in mouse and man and reach the smallest possible p-value (p<2.2×10^−16^).

### Defining the core T helper cell RBPome

To define a T helper cell RBPome we first made sure that RNA-IC and OOPS did not preferentially identify high abundance proteins (**Fig. 4a-b**). In comparison to single shot total proteomes OOPS-identified RBPs spanned the whole range of protein expression without apparent bias. In general, this was also true for RNA-IC, with a slight tendency to more abundantly expressed proteins. This however might be a true effect since messenger RNA binding RBPs have been reported to be higher in expression even in comparison to other RBPs ^22^. We used the recently established comprehensive eukaryotic RBP database as reference to compare OOPS and RNA-IC-identified canonical and non-canonical RBPs from mouse and human CD4^+^ T cells. The numbers of proteins in the mouse T helper cell RBPomes created by RNA-IC and OOPS ranging from 312 to 1255 made up 10% to 40% of all listed EuRBPDB proteins, respectively (**Fig. 4c**). OOPS-identified T cell RBPs outnumbered those from RNA-IC experiments by a factor of four, which was predominantly due to the eight times higher number of non-canonical RBPs. Interestingly though, there were also twice as many canonical RBPs significantly enriched by OOPS (**Fig. 4c**). Analyzing the ten most abundantly annotated mouse RBDs (comprising 26 to 224 members) showed that RNA-IC and OOPS often identified the same canonical RBPs (**Supplementary Fig. 5**), however at least equal, higher or much higher numbers were detected in OOPS samples depending on the specific RBD (**Fig. 4d**). These findings suggested that the OOPS method recovered RBPs from additional, non-polyadenylated RNAs and that RNA-IC-derived RBPomes are specific but incomplete. These conclusions are also supported by highly similar results obtained for the human CD4^+^ T cell RBPome (**Fig. 4e-f**). In a 4-way comparison of mouse and human RBPs identified by OOPS and RNA-IC (**Fig. 4g**) we conservatively defined all proteins that were identified by at least two datasets as ‘core CD4^+^ T cell RBPomes’ discovering 798 mouse and 801 human RBPs in this category (**Supplementary Table 1**). A sizable number of 519 mouse and 424 human proteins were exclusively enriched by the OOPS method. While we cannot rule out false-positives, more than 55% of the proteins of both subsets matched to EuRBPDB-listed annotations (not shown). These findings suggested that genuine RBPs are found even outside of the intersecting set of OOPs and RNA-IC identified proteins and that the definition of RBPomes profits from employing different biochemical approaches.

**Figure 4:**
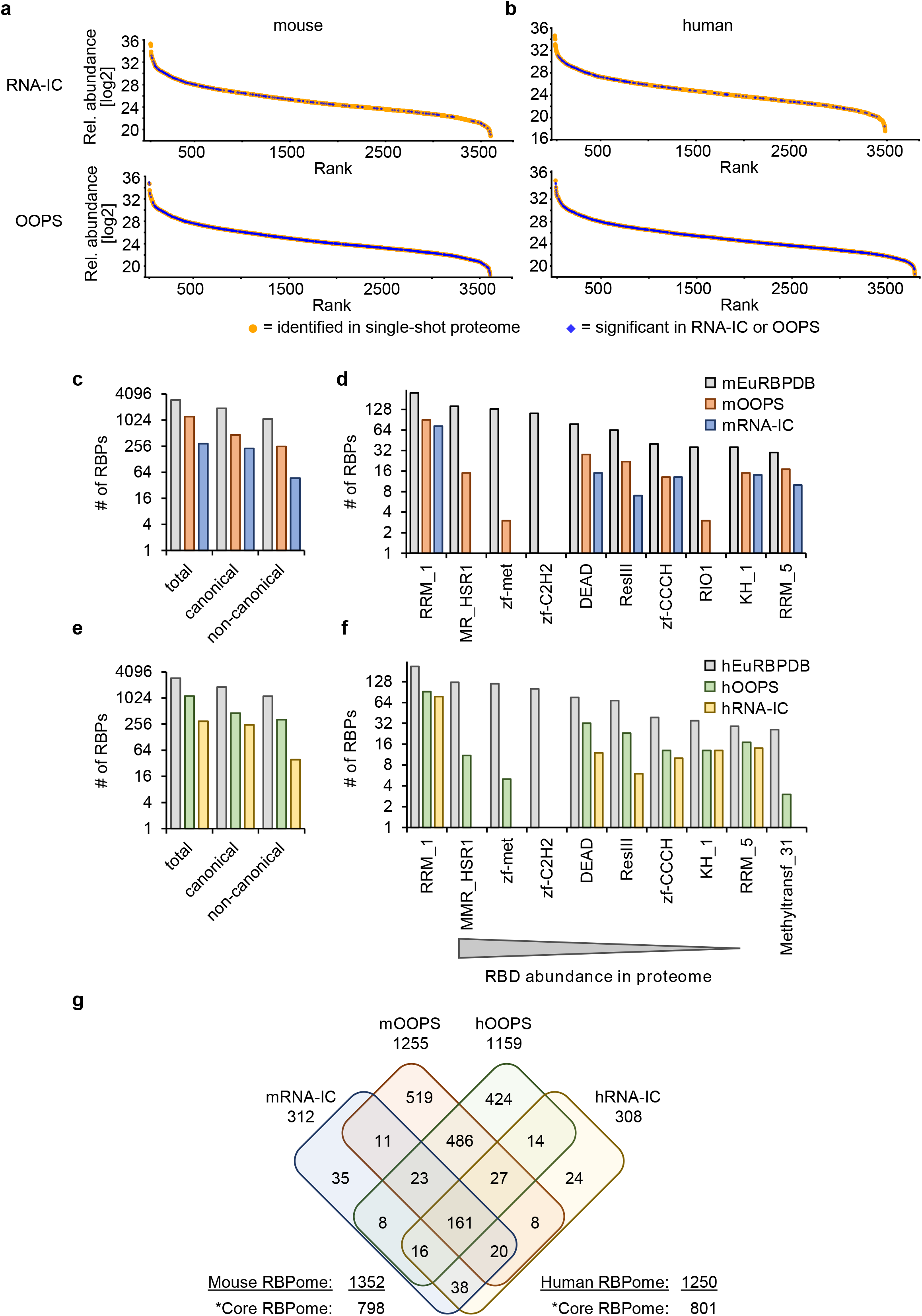
Defining the mouse and human CD4^+^ T cell RBPomes. **a,** Relative abundance (Log2) of proteins identified in a single-shot mouse proteome (orange dots) plotted by their rank from highest to lowest abundant protein. RNA-binding proteins detected by RNA capture (top plots) or by OOPS (bottom plots) are highlighted as blue diamonds. **b,** Same plots as shown in (a) for human data. **c, e,** The recently established database EuRBPDB was used as a reference for eukaryotic RBPs to determine the numbers of canonical and non-canonical RBPs identified by RNA-IC or OOPS on mouse (c) or human (e) CD4^+^ T cells. **d, f,** The occurrence of RBDs in RNA-IC and OOPS-identified RBPs was analyzed in comparison with the ten most abundant motifs described for the mouse (d) or human (f) proteome. **g,** Venn diagram using four datasets for RBPs in CD4^+^ T cells as determined by RNA-IC and OOPS in mouse and human cells. *The core RBPomes contain proteins present in at least two datasets.

### T cell signaling proteins with unexpected RNA-binding function

Some of the identified RBPs of the core proteome including Stat1, Stat4 and Crip1 are not expected to be associated with mRNA in cells. We therefore established an assay to confirm RNA-binding of these candidates. To do so, GFP-tagged candidate proteins were overexpressed in HEK293T cells, which were UV crosslinked, and extracts were used for immunoprecipitations with GFP specific antibodies. By Western or North-Western blot analyses with oligo(dT) probes we could verify pull-down of GFP-tagged proteins and the association with mRNA for the RBPs Roquin-1 and Rbms1, but also for the lactate dehydrogenase (Ldha) protein, a metabolic enzyme with known ability to also bind RNA ^23^ (**Fig. 5a**). Via this approach, the determined RNA association of Stat1, Stat4 and Crip1 by global methods was indeed confirmed (**Fig. 5b**), but appeared less pronounced as compared to prototypic RPBs, and similar with regard to Ldha (**Fig 5a**). This finding supports a potential moonlighting function of these signaling proteins. As the identity of the associated RNA species for these RBPs is unknown, we established a dual luciferase assay to determine the impact of the different proteins on the expression of the renilla luciferase transcript when they were tethered to an artificial 3*’*-UTR (**Fig. 5c**). We utilized the λN/5xboxB system ^41^ and confirmed the expression of fusion proteins with a newly established λN-specific antibody (**Fig. 5d-e**). Importantly, Stat1 and Stat4 repressed luciferase function almost to the same extend as the known negative regulators Pat1b and Roquin-1, or other known RBPs, such as Celf1, Rbms1 and Cpeb4 (**Fig. 5f**). λN-Crip1 and λN-Vav1 expression did not exert a positive or negative influence, since their relative luciferase expression appeared unchanged compared to cells transfected to only express the λN polypeptide (**Fig. 5f**). These data suggest that the signaling proteins Stat1 and Stat4 that we defined here as part of the T helper cell RBPome not only have the capacity to bind mRNA but can also exert RNA regulatory functions.

**Figure 5:**
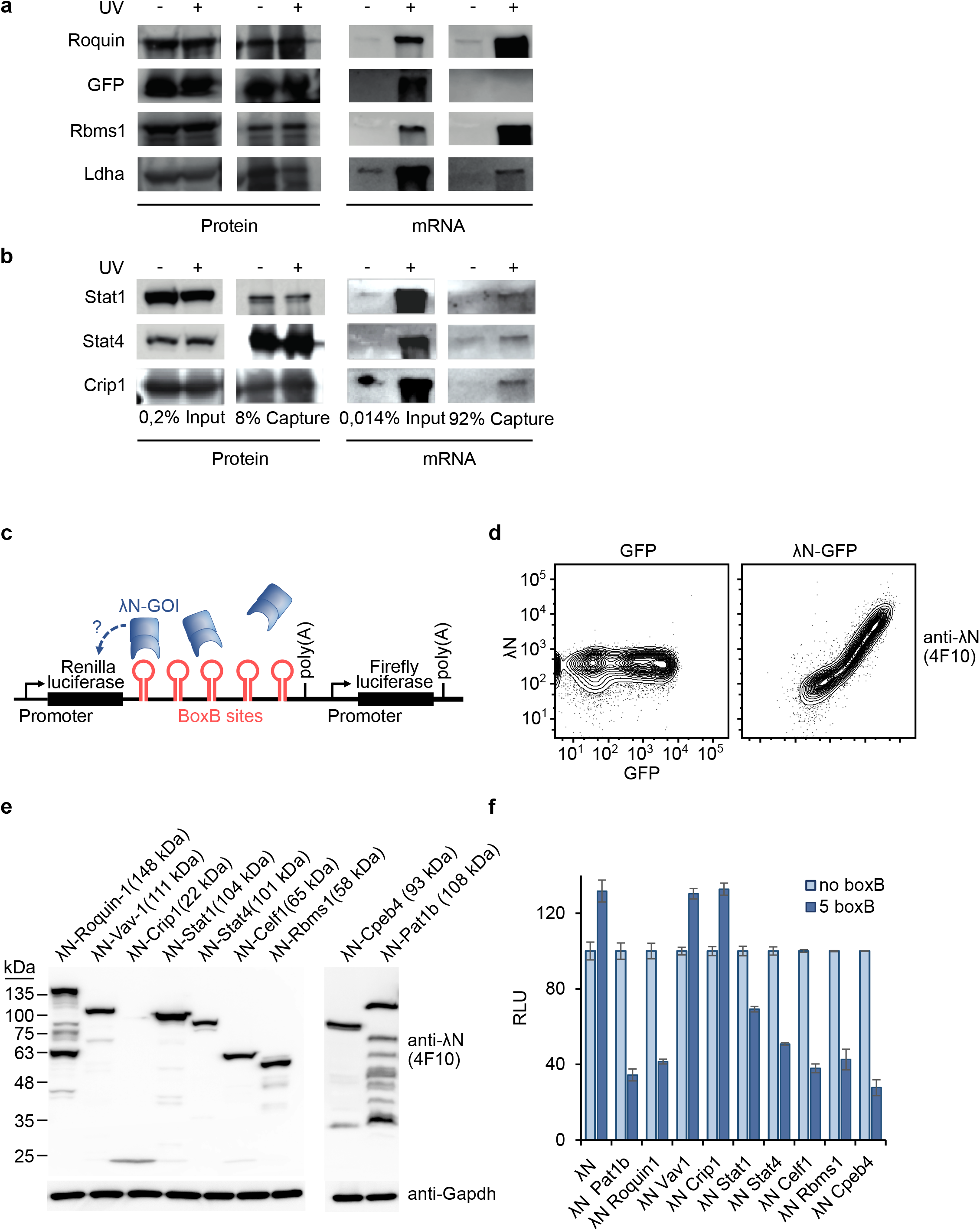
Stat1 and Stat4 are RBPs with regulatory potential. **a, b,** Semi-quantitative identification method for RBPs as in ^59^. In short, GFP-fused proteins were transfected into HEK293T cells, crosslinked or left untreated and subsequently immunoprecipitated using an anti-GFP antibody. The obtained samples were divided for protein and RNA detection by Western or Northern blotting, respectively. **c,** Schematic representation of the tethering assay that was used to investigate a possible influence of the genes of interest (GOI) on renilla luciferase expression. The affinity of the λN peptide targets the respective fusion protein to boxB stem-loop structures (5x) in the 3’-UTR of a Renilla luciferase gene where it can exert its function, if exciting. **d,** FACS plots using the new monoclonal antibody 4F10 demonstrate specific λN detection of a λN-GFP fusion protein expressed in 293T cells. **e,** Western blot showing the expression of the indicated λN-GOI proteins in 293T cells after transfection. **f,** Tethering assay results performed in HeLa cells as explained in (c). Two negative controls were implemented, using constructs without boxB sites or λN expression without fusion to a GOI. Each measurement was performed in triplicate and was independently repeated at least twice (n=2).

### Analyzing higher order post-transcriptional regulation

We next devised an experimental strategy to uncover RBPs with the potential to antagonize or cooperate with the Roquin-1 RBP in the repression of its target mRNAs by performing ‘BioID’ experiments (**Fig. 6a**). In this protein-centric, proximity-based labelling method we expressed a Roquin-1 BirA* fusion protein to identify proteins that reside within a short distance of approximately ∼10 nm ^42^ in T cells (**Fig. 6a**). In this dataset we sought for matches with the T cell RBPome (**Fig. 4g**) to identify proteins that shared the features, ‘RNA-binding’ and ‘Roquin-1 proximity’. We first verified that the mutated version of the biotin ligase derived from *E. coli* (BirA*) which was N-terminally fused to Roquin-1 or GFP was able to biotinylate residues in Roquin or other cellular proteins (**Fig. 6b**) but did not interfere with the ability of Roquin-1 to downregulate Icos (**Fig. 6c**). Doxycycline-induced BirA*- Roquin-1 compared to BirA*-GFP expression in CD4^+^ T cells significantly enriched biotin labelling of 64 proteins (**Supplementary Table 2**), including Roquin-1 (Rc3h1) itself or Roquin-2 (Rc3h2) (**Fig. 6d**) as well as previously identified Roquin-1 interactors and downstream effectors, such as Ddx6 and Edc4 ^26^, components of the Ccr4/Not complex ^43, 44^ and Nufip2 ^19^ (**Fig. 6d**). More than half of all proteins in proximity to Roquin-1 were also part of the defined RBPome (**Fig. 6e-f**). Increasing this BioID list with additional proteins that we determined in proximity to Roquin-1 when establishing and validating the BioID method in fibroblasts (**Supplementary Fig. 6a-d**), we arrived at 143 proteins (**Supplementary Table 2**) of which 96 (67%) were part of the RBPome (**Supplementary Fig. 6e** and **Supplementary Fig. 7a**). Cloning of 46 candidate genes in the context of N-terminal GFP fusions (**Supplementary Fig. 8a**) and generating retroviruses to overexpress candidate proteins we analyzed their effect on endogenous Roquin-1 targets (**Supplementary Fig. 8b**). CD4^+^ T cells were used from mice with *Rc3h1*^fl/fl^;*Rc3h2*^fl/fl^;rtTA alleles in combination with (iDKO) or without the CD4Cre-ERT2 allele (WT) allowing induced inactivation of Roquin-1 and −2 by 4’-OH-tamoxifen treatment. Reflecting deletion, the Roquin-1 targets Icos, Ox40, Ctla4, IκB_NS_ and Regnase-1 became strongly derepressed in induced double-knockout (iDKO) T cells (**Supplementary Fig. 8c**). This elevated expression was corrected to wildtype levels in iDKO T cells that were retrovirally transduced and doxycycline-induced to express GFP-Roquin-1 (**Supplementary Fig. 8c**). The target expression in WT T cells was only moderately reduced through overexpression of GFP-Roquin-1 (**Supplementary Fig. 8c**). For the majority of the 46 candidate genes induced expression in WT and iDKO CD4^+^ T cells did not alter expression of the five analyzed Roquin-1 targets, exemplified here by the results obtained for Vav1 (**Fig. 7a-b**). Interestingly, we identified a new function for Rbms1 (transcript variant 2), specifically upregulating Ctla-4 (**Fig. 7c-d**). Furthermore, we demonstrated that Cpeb4 strongly upregulates Ox40 and, most strikingly, in the same cells Cpeb4 repressed Ctla-4 levels (**Fig. 7e-f**). While these findings are new and noteworthy, they occurred in a Roquin-1 independent manner. In contrast to these effects, we determined a higher order regulation of Icos by Celf1. Induced Celf1 expression in wildtype T cells clearly upregulated Icos, Ctla-4 and Ox40 expression and this function was obliterated in Roquin-1-deficient iDKO cells (**Fig. 7g-h**). Importantly, this effect could not be explained by Celf1-mediated repression of Roquin-1 on the protein or mRNA level (**Supplementary Fig. 8d-e**). Together these findings demonstrate responsiveness of Roquin targets to many other RBPs and a higher order, Roquin-dependent regulation of transcripts encoding for the costimulatory receptors Icos, Ox40 and Ctla-4 by Celf1.

**Figure 6:**
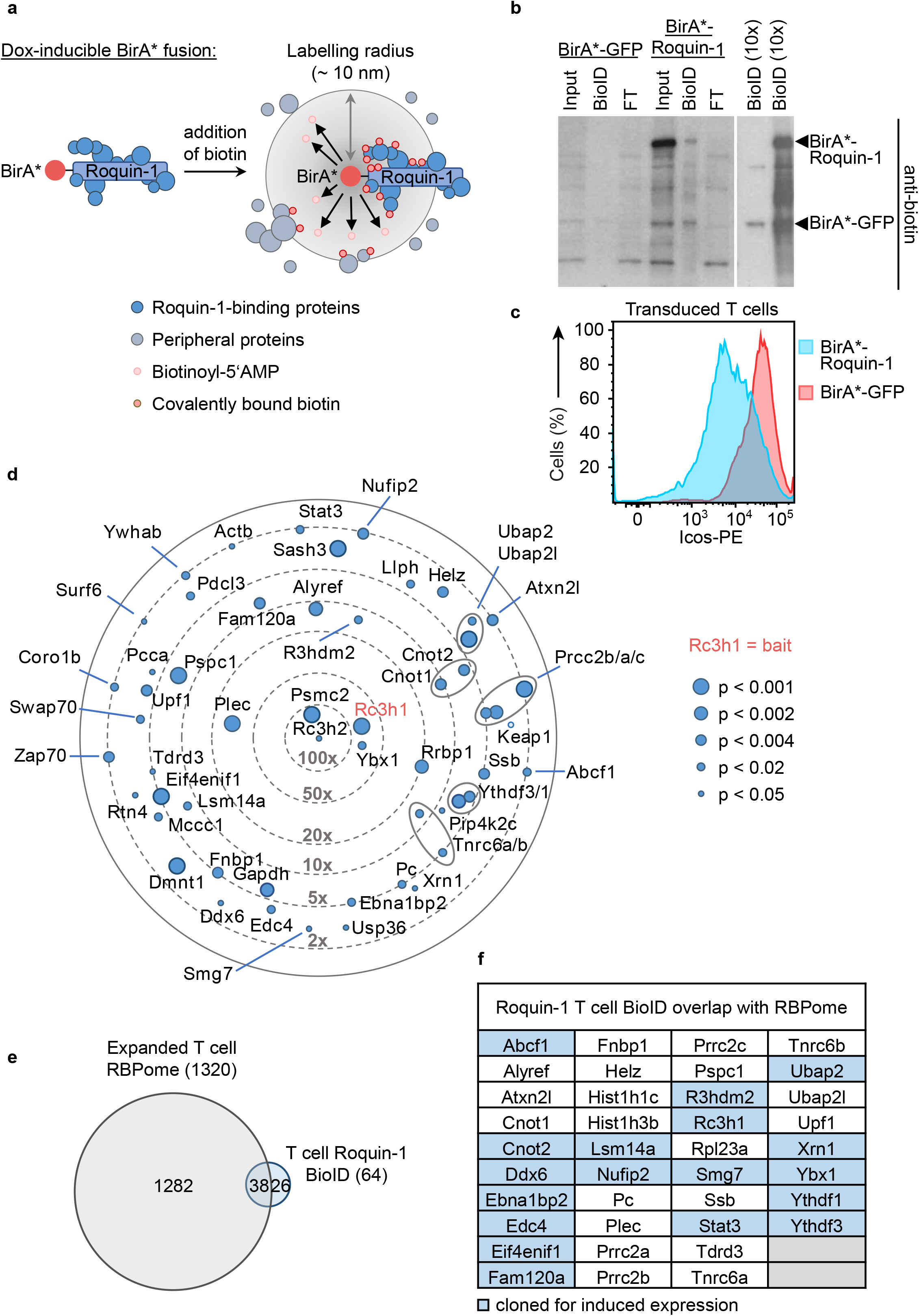
Identification of proteins in proximity to Roquin-1 in CD4^+^ T cells. **a,** Schematic overview of the BioID method showing how addition of biotin to the medium leads to the activation of biotin, diffusion of biotinoyl-5’AMP and the biotinylation of the bait (Roquin-1) and all preys in the circumference. **b,** Equimolar amounts of protein were loaded onto a PAGE gel for Western blotting applying an anti-biotin antibody. Efficient biotinylation of both baits BirA*-Roquin-1 and BirA*-GFP (control) could be demonstrated. **c,** Histogram showing that transduction of CD4^+^ T cells with retrovirus to inducibly express BirA*-Roquin-1 lead to the efficient downregulation of endogenous Icos. **d,** Identified preys from Roquin-1 BioIDs (n=5) in CD4^+^ T cells. Depicted are all significantly enriched proteins with the exception of highly abundant ribosomal and histone proteins. Dot sizes equal p-values and positioning towards the center implies increased x-fold enrichment over BirA*-GFP BioID results. **e,** Venn diagram showing the overlap of RBPs from the CD4^+^ T cell RBPome with the proteins identified by Roquin-1 BioID in T cells. **f,** Listed are all 38 proteins (59%) that are Roquin-1 preys and RBPs. Colored fields indicate proteins that were chosen for further analysis.

**Figure 7:**
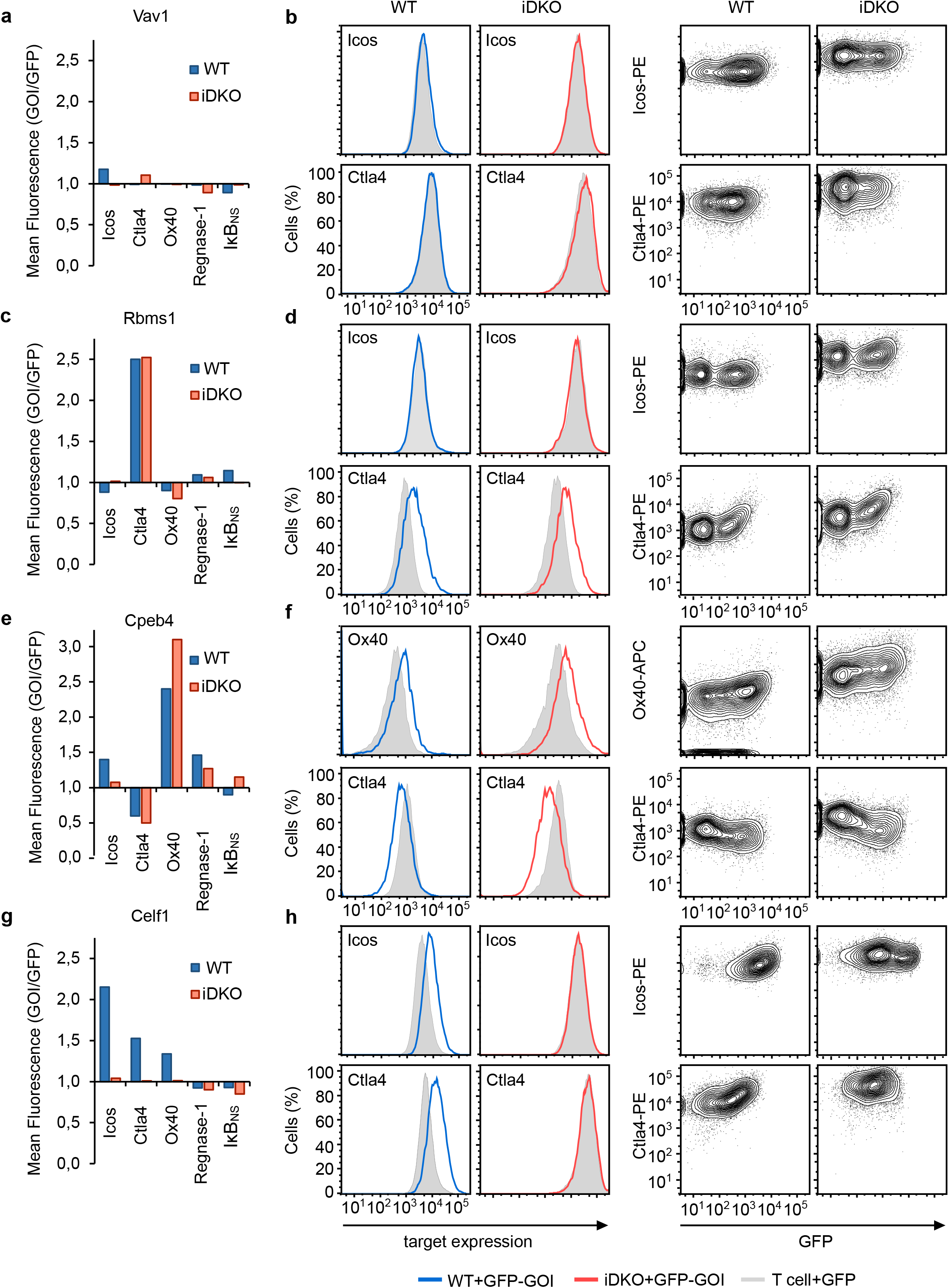
Higher order Icos regulation by Roquin-1 and Celf1. Treatment with 4’-OH-tamoxifen of CD4^+^ T cell with the genotypes Rc3h1^fl/fl^;Rc3h2^fl/fl^;rtTA3 without (WT) or with the CD4Cre-ERT2 allele (iDKO) were used for transduction with retroviruses. Expression levels of Icos and four additional Roquin-1 targets in WT and iDKO cells were analyzed 16h after individual, doxycycline-induced overexpression of 46 GFP-GOI fusion genes. **a, c, e, g,** The geometrical mean of each GOI-GFP divided by GFP for each Roquin-1 target was calculated and the summarized results are shown as bar diagrams for (a) Vav1, (b) Rbms1, (c) Cpeb4 and (d) Celf1. **b, d, f, h,** Representative, original FACS data are depicted as histograms or contour plots. Experiments for the negative Vav1 result were repeated twice, those for Rbms1, Cpeb4 and Celf1 at least three times.

## Discussion

The work on post-transcriptional gene regulation in T helper cells has focused on some miRNAs and several RNA-binding proteins, and few reports described m6A RNA methylation in this cell type. Although arriving at a more or less detailed understanding of individual molecular relationships and regulatory circuits, this isolated knowledge assembles into a very incomplete picture. Creating an atlas of the human and mouse T helper cell RBPomes has now opened the stage, allowing to work towards understanding connections, complexity and principles of post-transcriptional regulatory networks in these cells.

RNA-IC and OOPS are two complementary methods to define RBPs on a global scale. While RNA-IC specifically queries for proteins bound to polyadenylated RNAs, OOPS captures the RNA bound proteome in its whole. Applying both methods to T helper cells of two different organisms allowed us to cross-validate the results from both methods and solidified our description of the core mouse and human T helper cell RBPome. While the vast majority of RNA-IC-identified CD4^+^ T cell RBPs were previously known RNA binders, OOPS typically repeated and profoundly expanded these results (**Supplementary Fig. 5**), with the possible caveat of identifying false-positive proteins. Contradicting this possibility at least in part, more than half of OOPS identified proteins that were exclusively found in mouse or man were EuRBPDB-listed.

Strikingly, the signaling proteins Stat1 and Stat4 were identified by mouse RNA-IC and human OOPS and were just below the cut-off in the mouse OOPS dataset, and we could support their RBP function by additional RNA-binding assays. Undoubtedly, the defined human and mouse T cell RBPomes contain many more unusual RBPs that would warrant further investigations. We assume that even interactions of proteins without prototypic RBDs, like the Stat proteins, with RNA will have consequences for both binding partners. As the identity of the interacting mRNA(s) is unknown, we could only speculate about the post-transcriptional impact. Nevertheless, Stat1 and Stat4 showed regulatory capacity in our tethering assays. Intriguingly, RNA binding may also impact the function of Stat proteins as transcription factors. Supporting this notion, early results found Stat1 bound to the non-coding, polyadenylated RNA ‘TSU’, derived from a trophoblast cDNA library, and translocation of Stat1 into the nucleus was reduced after TSU RNA microinjection into HeLa cells ^45, 46^.

Many 3’-UTRs, which effectively instruct post-transcriptional control, exhibit little sequence conservation between species, and the exact modules which determine regulation are not known. This for example is true for the *Icos* mRNA ^19, 26^. On the side of the *trans*-acting factors, we find high similarity between the RBPomes of T helper cells of mouse and human origin, actually reflecting the general overlap of so far determined RBPomes from many cell lines of these species. We interpret this evolutionary conservation as holding ready similar sets of RBPs in T helper cells across species, which are then able to work together on composite *cis*-elements to reach comparable regulation of 3*’*-UTRs in the different organisms, despite sequence variability and differences in the composition of elements.

We not only define the first RBPomes of human and mouse T helper cells. We also provide potential avenues of how to make use of this information. Screening a set of candidates from the T cell RBPome for effects on Roquin targets, our findings support a concept in which post-transcriptional targets are separated into “regulons” ^47^. These regulons comprise coregulated mRNA subsets responding to the same inputs and often functioning in the same biological process. Therefore, complex and differential binding of targets by RBPs of the cellular RBPome specifies the possible regulons and differential functions of the cell. Roquin cooperated, coregulated, or antagonized in the regulation of Icos with Regnase-1, m6A methylation and miRNA functions. Moreover, our screening approach added Rbms1 as Roquin-independent and Celf1 as Roquin-dependent coregulator in the Icos containing regulon. It also revealed that different targets responded very differently to the expression of specific coregulators, as for example Ctla4 and Ox40 were inhibited or induced by the same RBP, Cpeb4, respectively. Together these data indicated an unexpected wealth of possible inputs from the T helper cell RBPome and suggested highly variable and combinatorial mRNP compositions in higher order post-transcriptional gene regulation of individual targets.

To solve the seemingly simple question which RBPs regulate which mRNA in T helper cells, we will require further knowledge about individual contributions, binding sites and composite *cis*-elements, temporal and interdependent occupancies, interactions among RBPs and with downstream effector molecules. In this endeavor global protein and RNA centric approaches make fundamental contributions.

## Acknowledgements

We thank Hemalatha Mutiah (Ludwig-Maximilians-Universität Munich) for screening hybridoma supernatants and also Claudia Keplinger (Helmholtz Center Munich) for excellent technical support. For the provision of mouse lines we would like to thank Marc Schmidt-Supprian (Rc3h1), Wolfgang Wurst (Rc3h2), Robert Blelloch (Dgcr8flox), Mingui Fu (Zc3h12aflox) and Thorsten Buch (Cd4-Cre-ERT2). The work was supported by the German Research Foundation grants SPP-1935 (to VH), SFB-1054 projects A03 and Z02 as well as HE3359/7-1 and HE3359/8-1 to V.H. as well as grants from the Wilhelm Sander, Fritz Thyssen, Else Kro□ner-Fresenius and Deusche Krebshilfe foundations to V.H‥ D.B. was supported by the Deutsche Forschungsgemeinschaft (DFG, German Research Foundation) under Emmy Noether Programme BA 5132/1-1 and BA 5132/1-2 (252623821), as well as SFB 1054 project B12 (210592381). D.B. is a member of Germany’s Excellence Strategy EXC2151 577 (390873048).

## Author contribution

EG and VH conceived the idea for the project and together with MW and MM supervised the experimental work. KPH and VH wrote the manuscript with contributions from EG, MW and AR. KPH performed OOPS, BioID and T cell transductions with help from GB, KD, SMH, JM and CC. CG conducted RNA-IC experiments and RNA-binding assays. AR and SMH performed mass spectrometry and AR analyzed RBPome data. SMH and KPH analyzed the BioID data. MX contributed the tethering assays and EW analyzed Icos regulation for which AG, RF, TI-K and DB provided unpublished reagents.

## Data availability statement

The data sets generated and/or analyzed during the current study are available from the corresponding authors on reasonable request.

## Methods

### Isolation, *in vitro* cultivation and transduction of primary CD4^+^ T cells

For *in vitro* cultivation of primary murine CD4^+^ T cells, mice were sacrificed and spleen as well as cervical, axillary, brachial, inguinal and mesenteric lymph nodes were dissected and pooled. Organs were squished and passed through a 100 μm filter under rinsing with T cell isolation buffer (PBS supplemented with 2% FCS and 1 mM EDTA). Erythrocytes were eliminated by incubating cells with TAC-lysis buffer (13 mM Tris, 140 mM NH_4_Cl, pH 7.2) for 5 min at room temperature. CD4^+^ T cells were isolated by negative selection using EasySep™ Mouse CD4^+^ T cell isolation Kit (STEMCELL) according to manufacturer’s protocol. Purified CD4^+^ T cells were cultured in DMEM (Invitrogen) T cell culture medium supplemented with 10% FCS (Gibco), 1% Pen-Strep (Thermo Fisher), 1% HEPES-Buffer (Invitrogen), 1% non-essential amino acids (NEAA; Invitrogen) and 1 mM β-mercaptoethanol (Invitrogen). For activation and differentiation under T_H_1 conditions the T cells were stimulated with α-CD3 (0.5 μg/mL; cl. 2C11, in house production), α-CD28 (2.5 μg/mL; cl. 37.5N, in house production), 10 μg/mL α-IL-4 (cl. 11B11, in house production) and 10 ng/mL IL-12 (BD Pharmingen) and cultured on goat α-hamster IgG (MP Biochemicals) pre-coated 6-or 12-well culture plates for 40-48h at an initial cell density of 5 or 1.5×10^6^ cells/mL, respectively. The cells were then resuspended and cultured in medium supplemented with 200 IE/mL recombinant hIL-2 (Novartis) in a 10% CO_2_ incubator and expanded for 2-4 days, as indicated. Subsequently cells were fed with fresh IL-2-containing medium every 24h and cultured at a density of 0.5-1×10^6^ cells/mL. For *in vitro* deletion of floxed alleles of Rc3h1^fl/fl^;Rc3h2^fl/fl^;CD4Cre-ERT2;rtTA3 (but with no effect on the Cre-deficient WT control Rc3h1^fl/fl^;Rc3h2^fl/fl^;rtTA3) CD4^+^ T cells were treated with 1 μM 4′OH-tamoxifen (Sigma) for 24h prior to activation at a cell density of 0,5-1×10^6^ cells/mL. We performed retroviral transduction 40h after the start of anti-CD3/CD28 activation, T cells were transduced with retroviral particles using spin-inoculation (1h, 18 °C, 850 g), and after 4-6h co-incubation of T cells and virus, virus particles were removed and T cells resuspended in T cell medium supplemented with IL-2 as described before. To induce expression of pRetro-Xtight-GFP constructs in rtTA expressing T cells, the transduced cells were cultured for 16h in the presence of doxycycline (1 μg/mL) prior to flow cytometry analysis of expression of targets in transduced GFP+ cells.

### *In vivo* deletion of loxP-flanked alleles and *in vitro* culture of CD4^+^ T cells

For *in vitro* culture analysis, deletion of Roquin (Rc3h1^fl/fl^;Rc3h2^fl/fl^;CD4-Cre-ERT2) ^48^, Regnase-1 (Zc3h12a^fl/fl^;CD4-Cre-ERT2) ^49^, Wtap (Wtap^fl/fl^;CD4-Cre-ERT2) ^33^, and Dgcr8 (Dgcr8^fl/fl^;CD4-Cre-ERT2) ^34^ encoding alleles in Cd4-cre-ERT2 mice was induced *in vivo* by oral transfer of 5 mg tamoxifen (Sigma) in corn oil. Two doses of tamoxifen each day were given on two consecutive days (total of 20 mg tamoxifen per mouse). Mice with the genotype CD4-Cre-ERT2 (without floxed alleles) were used for wild-type controls. Mice were scarified three days after the last gavage and total CD4^+^ T cells were isolated using the EasySep™ Mouse T Cell Isolation Kit (Stem Cell) and activated under T_H_1 conditions as described above.

### Flow cytometry

Following *in vivo* deletion and CD4^+^ T cell isolation (above) cells were activated and cultured under T_H_1 conditions. Cells were obtained daily for FACS analysis. The single cell suspensions were stained with fixable violet dead cell stain (Thermo Fisher) for 20 min at 4°C. For the detection of surface proteins, cells were stained with the appropriate antibodies in FACS buffer for 20 min at 4°C. After staining, cells were acquired on a FACS Canto II (3-laser). The data were further processed with the software FlowJo 10 software (BD Bioscience). The following antibodies were used: anti-CD4 (cl. GK1.5), anti-CD44 (cl. IM7), anti-CD62L (cl. MEL-14), anti-Icos (cl. C398.4A), anti-Ox40 (cl. OX-86), all from eBioscience, anti-CD25 (cl. PC61, Biolegend).

Effects of doxycycline-induced expression of 46 GFP-GOI fusion protein in 2×10^6^ wild-type and Roquin iDKO CD4^+^ T cells were analyzed on day 6 after isolation (compare **Suppl. Fig. 8b**). First, proteins were treated with a fixable blue dead cell stain (Invitrogen) and after washing, stained in three panels to interrogate the surface expression of Icos and Ox40 (Icos-PE, clone 7E.17G9/Ox40-APC, clone OX-86; both eBioscience) and to intracellularly measure Ctla4, IκB_NS_ (Ctla4-PE, UC10-4B9; eBioscience/cl. 4C1 rat monoclonal; in house production) as well as Regnase-1 (cl. 15D11 rat monoclonal; in house production). For intracellular staining cells were fixed in 2% formaldehyde for 15 min at RT, permeabilized by washing in Saponin buffer and stained with the appropriate antibodies for 1h at 4 °C. After washing, anti-rat-AF647 antibody (cl. poly4054; Biolegend) was added for 30 min. After additional rounds of Saponin- and FACS buffer washing, acquisition was performed using a LSR Fortessa.

### Isolation and differentiation of effector and regulatory T cells for RNA-IC

Naïve CD4^+^ T cells were isolated by using Dyna- and Detachabeads (11445D and 12406D, Invitrogen) from spleens and mesenteric lymph nodes of 8-12 week old C57BL/6J mice. For iTreg culture, cells were additionally selected for CD62L^+^ with anti-CD62L (clone: Mel14) coated beads. All cells were then activated with plate-bound anti-CD3 (using first anti-hamster, 55397, Novartis, then anti-CD3 in solution: clone: 2C11H: 0.1 μg/mL) and soluble anti-CD28 (clone: 37N: 1 μg/mL) and cultured in RPMI medium (supplemented with 10% (vol/vol) FCS, β-mercaptoethanol (0.05 mM, Gibco), penicillin-streptomycin (100 U/ml, Gibco), Sodium Pyruvate (1 mM, Lonza), Non-Essential Amino Acids (1x, Gibco), MEM Vitamin Solution (1x, Gibco), Glutamax (1x, Gibco) and HEPES pH 7.2 (10 mM, Gibco)). For iTreg cell differentiation we additionally added the following cytokines and blocking antibodies: rmIL-2 and rmTGF-β (both: R&D Systems, 5 ng/ml), anti-IL-4 (clone: 11B11, 10 μg/ml) and anti-IFNγ (clone: Xmg-121, 10 μg/ml). All antibodies were obtained in collaboration with and from Regina Feederle (Helmholtz Center Munich). After differentiation for 36-48 h cells were expanded for 2-3 days. iTreg cells were cultured in RPMI and 2000 units Proleukin S (02238131, MP Biomedicals) and T_eff_ cells with 200 U Proleukin S. We only used iTreg cells for experiments if samples achieved at least 80% Foxp3 positive cells (00552300, Foxp3 Staining Kit, BD Bioscience).

EL-4 T cells were cultured in the same medium as primary T cells. HEK293T cells were cultured in DMEM (supplemented with 10% (vol/vol) FCS, penicillin-streptomycin (100 U/ml, Gibco) and Hepes, pH 7.2 (10 mM, Gibco)).

### RBPome capture

For RBPome capture for mass spectrometry, 20 × 10^6^ T_eff_ or iTreg cells were either lysed directly (nonirradiated, control) in 1 ml lysis buffer from the μMACS mRNA Isolation Kit (130-075-201, Miltenyi) or suspended in 1 ml PBS, dispensed on a 10 cm dish and UV irradiated at 0.2 J/cm^2^ at 254 nm for 1 min, washed with PBS, pelleted and subsequently lysed (UV irradiated) and mRNA was isolated from both samples with the μMACS mRNA Isolation Kit according to the manufacturer’s instructions. RNAs and crosslinked proteins were eluted with 70° C RNase-free H_2_O. For RBPome capture for Western blot analysis, 400 × 10^6^ EL-4 T cells were either lysed directly in 8 ml lysis buffer (nonirradiated, control) or suspended in 16 ml PBS, dispensed on sixteen 10 cm dishes and UV irradiated at 0.2 J/cm^2^ at 254 nm for 1 min, washed with PBS, pelleted and subsequently lysed in 8 ml lysis buffer (UV irradiated) and then mRNA was isolated from both samples with the μMACS mRNA Isolation Kit using 500 μl oligo(dT) beads per sample. Each sample was split and run over two M columns (130-042-801, Miltenyi) and each column was eluted with two times 100 μl RNase-free H_2_O. The eluate was concentrated in Amicon centrifugal filter units (UFC501008, Merck) to a final volume of ∼25 μl. 8 μl Lämmli buffer (4x) with 10% (vol/vol) β-mercaptoethanol was added and the sample boiled for 5 min at 95° C. For protein analysis, samples were either flash-frozen in liquid nitrogen for MS analysis or Lämmli loading dye was added for subsequent analysis by Western blotting or silver staining.

### OOPS

20-30 × 10^6^ mouse CD4^+^ T cells were isolated as described above and activated without bias. Cells were washed in PBS once and resuspended in 1 ml of cold PBS and transferred into one well of a six well dish. Floating on ice, the cells in the open 6 well plate were UV irradiated once at 0.4 J/cm^2^ and twice at 0.2 J/cm^2^ at 254 nm with shaking in-between sessions. After irradiation the 1ml of cell suspension was transferred into a FACS tube and the well was washed with another 1 ml of cold PBS which was also added to the FACS tube. After centrifugation the cell pellet was completely dissolved in 1 ml of Trizol (Sigmal). The remainder of the procedure was performed strictly according to ^39^ with the exception that we broke up and regenerated interphases in five successive rounds of phase partitioning, rather than three.

### Culture preparation of human CD4^+^ T cell blasts

Blood (120 ml) was collected by venepuncture from two times four donors for RNA-IC and OOPS. Peripheral blood mononuclear cells (PBMCs) were isolated by standard Ficoll gradient (Pancoll^R^) centrifugation and CD4^+^ T cells were isolated from 1×10^8^ cells using CD4^+^ Microbeads (Miltenyi) to arrive at 2-3×10^7^ cells. These were resuspended at 2×10^6^/ml in T cell medium ((AIM-V (Invitrogen), 10% heat-inactivated human serum, 2mM L-glutamine, 10 mM HEPES and 1,25 μg/ml Fungizone), supplemented with 500 ng/ml PHA (Murex) and 100 IU IL-2/ml (Novartis). Cells were dispensed in 24-well plates at 2 ml per well and on day 3 old medium was replaced by fresh T cell medium and cell were harvested and counted at day 4,5. For generating iTreg, naïve CD4^+^ T cells were isolated from PBMCs using Microbeads (Naive CD4+ T Cell Isolation Kit II, Miltenyi) and resuspended at 1×10^6^ cells/ml in T cell medium supplemented with 500 ng/ml PHA (Murex), 500 IU IL-2/ml (Novartis), 500 nM Rapamycin (Selleckchem), and 5 ng/ml TGF-ß1 (Miltenyi). Cells were cultured in 24-well plates at 2 ml per well together with 1×10^6^ irradiated (40 Gy) PBMCs derived from three donors. On day 3, old medium was replaced by fresh T cell medium including supplements and the cells harvested and counted at day 5.

### SDS-PAGE, Western blotting and silver staining

All protein visualization procedures were performed according to standard protocols. For silver staining we used the SilverQuest kit (LC6070, Invitrogen). Antibodies used were anti-GFP, (1:10, clone: 3E5-111, in house), anti-Ptbp1, (1:1000, 8776, Cell Signaling), anti-β- tubulin, (1:1000, 86298, Cell Signaling). Proteins were visualized by staining with anti-rat (1:3000, 7077, Cell Signaling) or anti-mouse (1:3000, 7076, Cell Signaling) secondary antibodies conjugated to HRP.

### Sample preparation for mass spectrometry

For RNA-capture, eluates were incubated with 10 μg/ml RNase A in 100 mM Tris, 50 mM NaCl, 1 mM EDTA at 37°C for 30 min. RNase-treated eluates were acetone precipitated and resuspended in denaturation buffer (6 M urea, 2 M thiourea, 10 mM Hepes, pH 8), reduced with 1 mM DTT and alkylated with 5.5 mM IAA. Samples were diluted 1:5 with 62.5 mM Tris, pH 8.1 and proteins digested with 0.5 μg Lys-C and 0.5 μg Trypsin at room temperature overnight. The resulting peptides were desalted using stage-tips containing three layers of C18 material (Empore).

For OOPS experiments, 100 μl of lysis buffer (Preomics, iST kit) were added and samples incubated at 100°C for 10 min at 1,400 rpm. Samples were sonicated for 15 cycles (30s on/30s off) on a bioruptor (Diagenode). Protein concentration was determined using the BCA assay and about 30 μg of proteins were digested. To this end, trypsin and Lys-C were added in a 1:100 ratio, samples diluted with lysis buffer to contain at least 50 μl of volume and incubated overnight at 37°C. To 50 μl of sample, 250 ul Isopropanol/1% TFA were added and samples vortexed for 15s. Samples were transferred on SDB-RPS (Empore) stagetips (3 layers), washed twice with 100 μl Isopropanol/1% TFA and twice with 100 μl 0.2% TFA. Peptides were eluted with 80 μl of 2% ammonia/80% acetonitrile, evaporated on a centrifugal evaporator and resuspended with 10 μl of buffer A* (2% ACN, 0.1% TFA).

### LC-MS/MS analysis

Peptides were separated on a reverse phase column (50 cm length, 75 μm inner diameter) packed in-house with ReproSil-Pur C18-AQ 1.9 μm resin (Dr. Maisch GmbH). Reverse-phase chromatography was performed with an EASY-nLC 1000 ultra-high pressure system, coupled to a Q-Exactive HF Mass Spectrometer (Thermo Scientific) for mouse RNA-capture experiments or a Q-Exactive HF-X Mass Spectrometer (Thermo Scientific) for human RNA-capture, OOPS experiments and single-shot proteomes. Peptides were loaded with buffer A (0.1% (v/v) formic acid) and eluted with a nonlinear 120-min (100-min gradient for human RNA-capture and OOPS experiments) gradient of 5–60% buffer B (0.1% (v/v) formic acid, 80% (v/v) acetonitrile) at a flow rate of 250 nl/min (300 nl/min for human RNA-capture and OOPS). After each gradient, the column was washed with 95% buffer B and re-equilibrated with buffer A. Column temperature was kept at 60° C by an in-house designed oven with a Peltier element and operational parameters were monitored in real time by the SprayQc software. MS data were acquired using a data-dependent top 15 (top 12 for human RNA-capture and OOPS experiments) method in positive mode. Target value for the full scan MS spectra was 3 × 10^6^ charges in the 300−1,650 m/z range with a maximum injection time of 20 ms and a resolution of 60,000. The precursor isolation windows was set to 1.4 m/z and capillary temperature was 250°C. Precursors were fragmented by higher-energy collisional dissociation (HCD) with a normalized collision energy (NCE) of 27. MS/MS scans were acquired at a resolution of 15,000 with an ion target value of 1 × 10^5^, a maximum injection time of 120 ms (60 ms for human RNA-capture and OOPS experiments). Repeated sequencing of peptides was minimized by a dynamic exclusion time of 20 s (30 ms for human RNA-capture and OOPS).

### Raw data processing

MS raw files were analyzed by the MaxQuant software^50^ (version 1.5.1.6 for RNA-capture files and version 1.5.6.7 for OOPS files) and peak lists were searched against the mouse or human Uniprot FASTA database, respectively, and a common contaminants database (247 entries) by the Andromeda search engine^51^. Cysteine carbamidomethylation was set as fixed modification, methionine oxidation and N-terminal protein acetylation as variable modifications. False discovery rate was 1% for both proteins and peptides (minimum length of 7 amino acids). The maximum number of missed cleavages allowed was 2. Maximal allowed precursor mass deviation for peptide identification was 4.5 ppm after time-dependent mass calibration and maximal fragment mass deviation was 20 ppm. Relative quantification was performed using the MaxLFQ algorithm^52^. “Match between runs” was activated with a retention time alignment window of 20 min and a match time window of 0.5 min for RNA-capture experiments, while matching between runs was disabled for OOPS experiments. The minimum ratio count was set to 2 for label-free quantification.

### Data analysis

Statistical analysis of MS data was performed using Perseus (version 1.6.0.28). Human RNA-capture, mouse RNA-capture, human OOPS and mouse OOPS data was processed separately. For all experiments, MaxQuant output tables were filtered to remove protein groups matching the reverse database, contaminants or proteins only observed with modified peptides. Next, protein groups were filtered to have at least two valid values in either the crosslinked or control triplicate. LFQ intensities were logarithmized (base 2) and missing values were imputed from a normal distribution with a downshift of 1.8 standard deviations and a width of 0.2 (0.25 for OOPS data). For RNA-capture experiments, a Student’s T-test was performed to find proteins significantly enriched in the crosslinked sample over the non-crosslinked control (false-discovery rate (FDR) < 0.05). As many proteins were identified specifically in the crosslinked sample at intensities too low to find significant differences compared to the imputed values, we additionally considered proteins only identified in two or three replicates of the crosslinked sample, but never in the non-crosslinked control, as RNA-binding proteins. For OOPS experiments, proteins significantly enriched in a Student’s T-tests of the organic phase after RNase-treatment over the same sample of the non-crosslinked control (FDR < 0.05) were considered RBPs.

GO term enrichment analysis was performed using the clusterProfiler package in R (version 4.1.0) as described in the original publication^53^. The mouse or human proteomes served as background for the respective enrichment analysis. Relative abundance of proteins in the single-shot proteome (**Fig. 4a-b**) was determined by calculating the logarithm (base 2) of the ratio of the LFQ intensity and the number of theoretical peptides. Relative abundance of proteins significant in the mouse OOPS and RNA-IC is shown in the single-shot proteome of mouse CD4^+^ T cells measured with the OOPS samples. Relative abundance of proteins significant in the human OOPS and RNA-IC experiments is shown in the single-shot proteome of human CD4^+^ T cells measured with the OOPS samples or RNA-IC samples, respectively. For the 4-way Venn comparison to find RBPs identified in more than one RNA-IC or OOPS experiment, human gene names were converted into their homologous mouse counterparts. Protein group entries containing more than one isoform were expanded for the comparison and subsequently collapsed into one entry again for calculation of the Venn diagram. Multiple gene name entries for different unambiguously identified isoforms were collapsed into the major isoform. This affected six protein groups in the human RNA-IC dataset and three protein groups in the mouse RNA-IC dataset.

Intrinsically disordered regions were retrieved from the Disorder Atlas^54^. We referred to the LCR-eXXXplorer^55^ to obtain low complexity regions in proteins.

### Plasmid construction

To generate a vector that expresses N-terminally GFP-tagged proteins, we amplified the respective genes from cDNA of T_eff_ or T_Foxp3+_ cells, added *Hind*III and *Kpn*I restriction sites in front of the start codon and cloned them into the pCR™8/GW/TOPO® vector. We then used *Hind*III and *Kpn*I to insert a GFP sequence where we removed the bases for the stop codon. The respective sequences were subsequently transferred to the expression vector pMSCV via the gateway cloning technology. Only GFP-Roquin-1 (a kind gift of Vigo Heissmeyer) was expressed from the vector pDEST14. For oligonucleotide sequences see **Supplementary Table 3**.

### Validation of RNA binding ability

HEK293T cells were transfected by calcium phosphate transfection with plasmids expressing the respective proteins with an N-terminal GFP-tag or GFP alone. After three days, cells were washed with PBS on plates, UV crosslinked (CL) as before or directly scraped from the plates (nCL). Cell lysates were generated by flash-freezing pellets in liquid nitrogen and incubation in NP-40 lysis buffer (150 mM NaCl, 1% NP-40, 50 mM Tris-HCl, pH 7.4, 5 mM EDTA, 1 mM DTT, 1 mM PMSF and protease inhibitor mixture (Complete, Roche)). After lysis, extracts were cleared by centrifugation at 17000 g for 15 min at 4° C. We then determined protein concentration via the BCA method and used 2-10 mg of protein for the subsequent GFP immunoprecipitation, depending on transfection efficiency and expected RNA-binding capacity. We pre-coupled 200 μl Protein-G beads (10004D, Dynabeads Protein G, Invitrogen) with 20 μg antibody (anti-GFP, clone: 3E5-111, in house) in PBS (1 h, RT), washed beads in lysis buffer, added them to cell lysates and incubated with rotation for 4 h at 4 °C. Beads were then washed three times with IP wash buffer (50 mM Tris-HCl, pH 7.5) with decreasing salt (500 mM, 350 mM, 150 mM, 50 mM NaCl) and SDS (0.05%, 0.035%, 0.015%, 0.005%) concentrations. Proteins and crosslinked RNAs were eluted with 50 mM glycine, pH 2.2 at 70° C for 5 min. Lämmli buffer (4x) was added and samples were divided for mRNA and protein detection and separated via SDS gel electrophoresis (6% SDS gels for detection of mRNA samples and 9% gels to verify immunoprecipitation efficiency). For RNA detection we blotted onto Nitrocellulose membranes and for protein detection on PVDF membranes. After transfer, the Nitrocellulose membrane was prehybridized with Church buffer (0.36 M Na_2_HPO_4_, 0.14 M NaH_2_PO_4_, 1 mM EDTA, 7% SDS) for 30 min and then incubated for 4 h with Church buffer containing 40 nM 3′-and 5′-Biotin labeled oligo(dT)_20_ probe to anneal to the poly-A tail of the bound mRNA. The membrane was washed twice with 1 x SSC, 0.5% SDS and twice with 0.5 x SSC, 0.5% SDS. Bound mRNA was detected with the Chemiluminescent Nucleic Acid Detection Kit Module (89880, Thermo Fisher) according to the manufacturer′s instructions.

### Real-Time PCR

Total RNA of input was purified from lysates with Agencourt RNAClean XP Beads (A63987, Beckman Coulter) according to the manufacturer’s instructions and eluted in nuclease-free H_2_O. cDNA was synthesized from total input RNA and oligo(dT)-isolated RNA with the QuantiTect Reverse Transcription Kit (205311, Qiagen). All qRT-PCRs were performed with the SYBR green method. For Primer sequences see **Supplementary Table 3**.

RNA isolation, reverse transcription and quantitative RT PCR as shown in Figure 8d was performed as published ^56^ using the universal probes systems (Roche). Primers for Rc3h1 (F: gagacagcaccttaccagca; R: gacaaagcgggacacacat; probe 22) and Hprt (F: Tgatagatccattcctatgactgtaga; R: aagacattctttccagttaaagttgag; probe 95) were efficiency tested (both E=1.99).

### Tethering assay

Hela cells were seeded in 24-well plates using 5×10^4^ cells per well. Transfection was performed the following day using Lipofectamine2000 (Invitrogen) and 300 ng of total constructs. Each transfection consisted of 75 ng of luciferase reporter plasmid psiCHECK2 (Promega) or luciferase-5boxB plasmid psiCHECK2 −5boxB, 225 ng of pDEST12.2-λN fused constructs. After 24 h, cells were harvested for luciferase activity assays using a Dual-Luciferase Reporter Assay System (Promega). Renilla luciferase activity was normalized to Firefly luciferase activity in each well to control for variation in transfection efficiency. psiCHECK2 lacking boxB sites served as a negative control, and each transfection was analysed in triplicates.

### BioID

The proximity-dependent biotin identification assay was performed according to Roux (Roux et al., 2012) with modifications. For each sample 2×10^7^ MEF cells were grown on ten 15-cm cell culture dishes for 24h before BirA*-Roquin-1 or BirA* expression was induced by doxycycline treatment. For T cells, transduction with the same BirA*-fusions cloned into the plasmid pRetroXtight was performed as described above and the same number of cells was used for the experiment. Six hours after addition of doxyclycline, biotin was added for 16h to arrive at an end concentration of 50 μM. Approximately 8×10^7^ cells per sample were trypsinized, washed twice with PBS and lysed in 5 ml lysis buffer (50 mM Tris-HCL, ph7,4; 500 mM NaCl, 0,2% SDS; 1x protease inhibitors (Roche), 20 mM DTT, 25 U/ml Benzonase) for 30 min at 4 °C using an end-over-end mixer. After adding 500 μl of 20% Triton X-100 the samples were sonicated for two sessions of 30 pulses at 30% duty cycle and output level 2, using a Branson Sonifier 450 device. Keep on ice for two minutes in between sessions.

Pipetting of 4,5 ml prechilled 50 mM Tris-HCL, pH 7,4 was followed by an additional round of sonication. During centrifugation at 16500 × g for 10 min at 4 °C 500 μl streptavidin beads (Invitrogen) or each sample was equilibrated in a 1:1 mixture of lysis buffer and 50 mM Tris-HCL pH 7,4. After overnight binding on a rotator at 4 °C Streptavidin beads were stringently washed using wash buffers 1, 2 and 3 (Roux et al, 2012) and prepared for mass spectrometry by three additional washes with buffer 4 (1 mM EDTA, 20 mM NaCl, 50 mM Tris-HCL pH 7,4). Proteins were eluted from streptavidin beads with 50 μl of biotin-saturated 1x sample buffer (50 mM Tris-HCL pH 6,8, 12% sucrose, 2% SDS, 20 mM DTT, 0,004% Bromphenol blue, 3 mM Biotin) by incubation for 7 min at 98 °C. For identification and quantification of proteins, samples were proteolysed by a modified filter aided sample preparation ^57^ and eluted peptides were analysed by LC-MSMS on a QExactive HF mass spectrometer (ThermoFisher Scientific) coupled directly to a UItimate 3000 RSLC nano-HPLC (Dionex). Label-free quantification was based on peptide intensities from extracted ion chromatograms and performed with the Progenesis QI software (Nonlinear Dynamics, Waters). Raw files were imported and after alignment, filtering and normalization, all MSMS spectra were exported and searched against the Swissprot mouse database (16772 sequences, Release 2016_02) using the Mascot search engine with 10 ppm peptide mass tolerance and 0.02 Da fragment mass tolerance, one missed cleavage allowed, and carbamidomethylation set as fixed modification, methionine oxidation and asparagine or glutamine deamidation allowed as variable modifications. A Mascot-integrated decoy database search calculated an average false discovery of < 1% when searches were performed with a mascot percolator score cut-off of 13 and significance threshold of 0.05. Peptide assignments were re-imported into the Progenesis QI software. For quantification, only unique peptides of an identified protein were included, and the total cumulative normalized abundance was calculated by summing the abundances of all peptides allocated to the respective protein. A t-test implemented in the Progenesis QI software comparing the normalized abundances of the individual proteins between groups was calculated and corrected for multiple testing resulting in q values (FDR adjusted p values) given in **Supplemental Table 2**. Only proteins identified by two or more peptides were included in the list of Roquin-1 proximal proteins.

### Antibodies

To generate monoclonal antibodies against pan-Ythdf proteins or λN-peptide, Wistar rats were immunized with purified GST-tagged full-length mouse Ythdf3 protein or ovalbumin-coupled peptide against λN (MNARTRRRERRAEKQWKAAN) using standard procedures as described ^58^. The hybridoma cells of Ythdf- or λN-reactive supernatants were cloned at least twice by limiting dilution. Experiments in this study were performed with anti-pan-Ythdf clone DF3 17F2 (rat IgG2a/κ) and anti-λN clone LAN 4F10 (rat IgG2b/κ).

---

**Supplementary Fig. 1: RNA-IC supporting results.**

**a,** Western blots demonstrating specificity of the newly established pan-Ythdf monoclonal antibody 17F2 for N-terminal GFP fusions of Ythdf1, Ythdf2 and Ythdf3, which are absent from non-transfected 293T cells. **b,** Silver staining analysis of oligo(dT) captured samples with and without UV irradiation. **c,** Quantitative RT-PCR to determine RNA pull-down efficiency with (CL) and without crosslink (nCL). Error bars show the standard deviation around the means of three independent experiments. **d,** Western blotting of UV irradiated and nonirradiated samples of EL-4 T cells. Membranes were probed with antibodies for the known mRNA-binding protein polypyrimidine tract binding protein 1 (Ptbp1) and β-tubulin. **e,** Distribution of low complexity regions in all Uniprot reviewed protein sequences (black line), in proteins in the EuRBPDB database (green) and in proteins significant in the RNA-IC data (red line). The left plot shows mouse data and the right plot human data. According to Kolmogorov-Smirnov testing the LCR distribution differences between RNA-IC (red lines) and all proteins (black lines) are highly significant in mouse (p<2.2×10^−16^) and man (p=7.8×10^−16^).

**Supplementary Fig. 2: RNA-IC on mouse and human iTreg cells. a,** Volcano plot showing the −log10 p-value in relation to the log2 fold-change comparing the RNA-capture from crosslinked mouse regulatory T cells (left plot) or human regulatory T cells (right plot) versus the non-crosslinked control. Red dots represent proteins significant at a 5% FDR cutoff level in both mouse and human RNA-capture experiments and blue dots proteins significant only in mouse or human, respectively. **b,** Venn diagram using four datasets to compare RNA-IC derived RBPomes of effector T and induced Treg cells from mouse and man.

**Supplementary Fig. 3: OOPS supporting results. a,** Agarose gel demonstrating disappearance of the typical 18S/28S rRNA bands after crosslinking and appearance of upshifted protein-RNA adducts (black arrows) in MEF cells with and without doxycycline-induce Roquin-1 expression. rRNA bands reappear after proteinase K (PK) treatment (grey arrows) and after RNase treatment the protein-RNA adducts in the wells disappear. **b,** Western blots showing that known RBPs, such as Roquin-1 and Gapdh can be detected in interphases, in the case of Roquin-1 only after induced expression. * cleavage product. **c, d,** Volcano plot showing the −log10 p-value plotted against the log2 fold-change comparing the interphase of OOPS experiments of crosslinked mouse CD4^+^ T cells versus the organic phase after RNase treatment of the interphase. Glycoproteins are highlighted in red. Enrichment analysis of GO Molecular Function terms was performed for proteins with a log2 fold-change larger than 2. The 20 most enriched terms are depicted.

**Supplementary Figure 4: Gene ontology enrichment analysis.** Enrichment analysis of GO Biological Process and GO Cellular Component terms of significant proteins in mouse or human RNA-IC data (top row) or OOPS data (bottom row). The ten most enriched terms in mouse (dark blue) and the respective terms in human (light blue) are shown. The y-axis represents the number of proteins matching the respective GO term. Numbers above each term depict the adjusted p-value after multiple testing correction (Benjamini-Hochberg).

**Supplementary Fig. 5: All currently annotated CCCH and KH domain containing RBPs and their detection in the RNA-IC and OOPS-identified RBPomes.** All RBPs with the indicated RBDs are listed according to EuRBPDB. Colored boxes indicate significant enrichment by RNA-IC or OOPS in mouse or human CD4^+^ T cells.

**Supplementary Fig. 6: Identification of Roquin-1 preys by BioID in MEF cells. a,** Western blots showing doxycycline-induced expression of Myc-BirA*-Roquin-1 or Myc-BirA* in MEF cell clones. **b,** FACS blot demonstrating that as in T cells (Fig. 6c) N-terminal fusion of Myc-BirA* to Roquin-1 does not affect its function and **c,** the fusion protein can biotinylate Roquin-1 and **d,** its cofactor Nufip2. **e,** Identified preys from Roquin-1 BioID in MEF cell clones. Depicted are 55 of 143 significantly enriched proteins (n=4) for better comparison with Fig. 6d. Yellow dot color indicates identification of the Roquin-1 prey in MEF and T cells. Dot sizes equal p-values and positioning towards the center implies increased x-fold enrichment.

**Supplementary Fig. 7: Overlap between Roquin-1 preys in MEF cells and the CD4^+^ T cell RBPome. a,** Venn diagram showing that 96 proteins (67%) are Roquin-1 preys and also RBPs in T cells. **b,** Table listing these 96 proteins. Colors indicate which genes were clone for downstream experiments and if they were identified by MEF BioID only (red) or additionally in the T cell BioID.(blue).

**Supplementary Fig. 8: Supporting results for higher order Icos regulation by Roquin-1 and Celf1. a,** Microscopical images showing different localizations of the GFP signal as a result of GFP-GOI subcellular targeting in transfected 293T cells. **b,** Schematic representation of the experiment performed in Fig. 7. **c,** Treatment with 4’-OH-tamoxifen of CD4^+^ T cell with the genotypes Rc3h1^fl/fl^;Rc3h2^fl/fl^;rtTA3 without (WT) or with the CD4Cre-ERT2 allele (iDKO) were used for transduction with a retrovirus expressing GFP-Roquin-1. Expression levels of the Roquin-1 targets Icos, Ox40, Ctla4 and Iκb_NS_ were analyzed and work as a positive control for the experiment. **d, e,** Cells taken from an experiment as performed (b). qPCR (d) and Western blot (e) performed on cDNA or protein lysates, respectively, derived from CD4^+^ T cells (WT) after doxycycline-induced expression of the indicated fusion proteins.

