## Supplementary figures and images for "Defining the RBPome of T helper cells to study higher order post-transcriptional gene regulation"

### Supplementary Figure 1

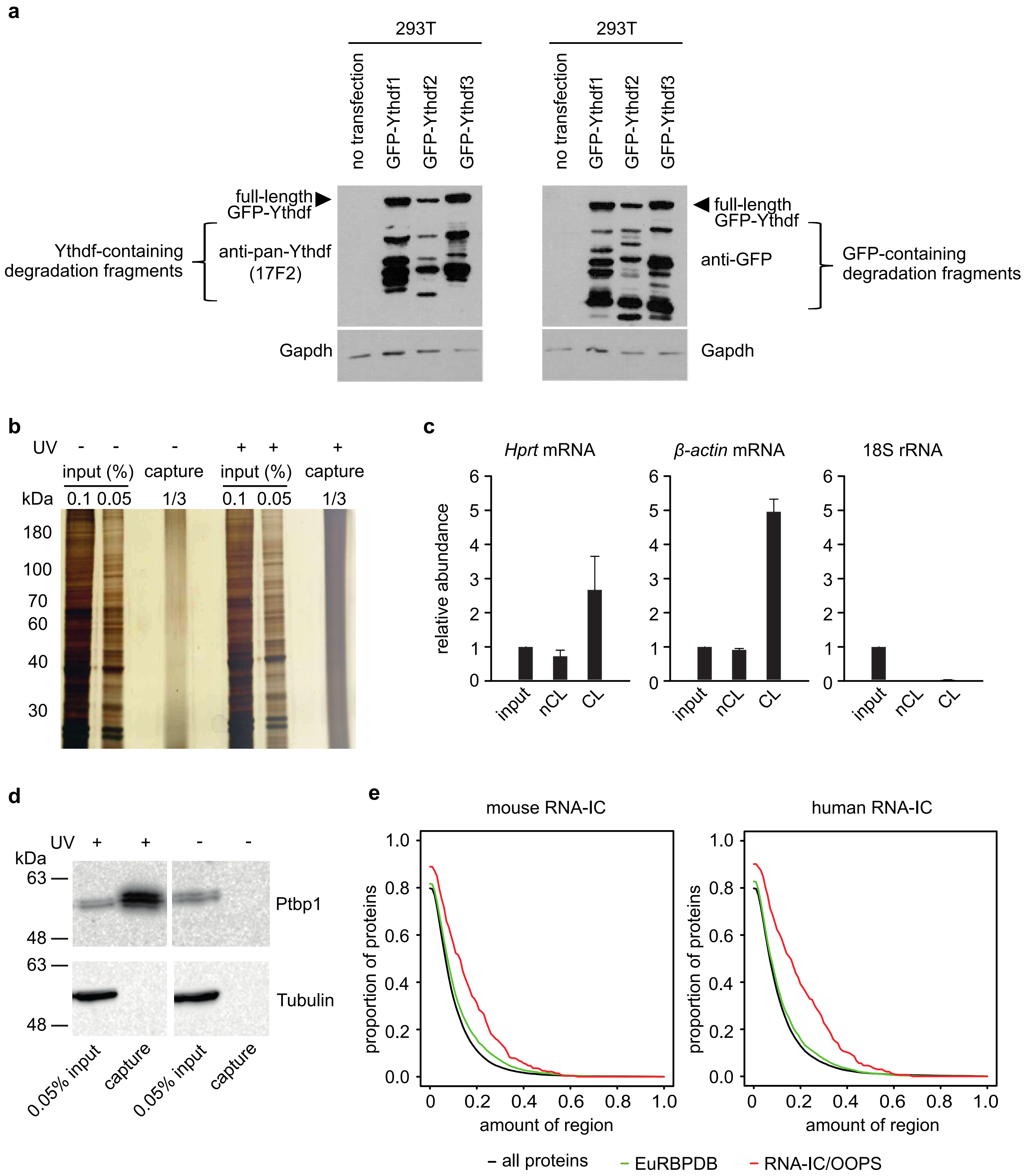

### Supplementary Figure 2

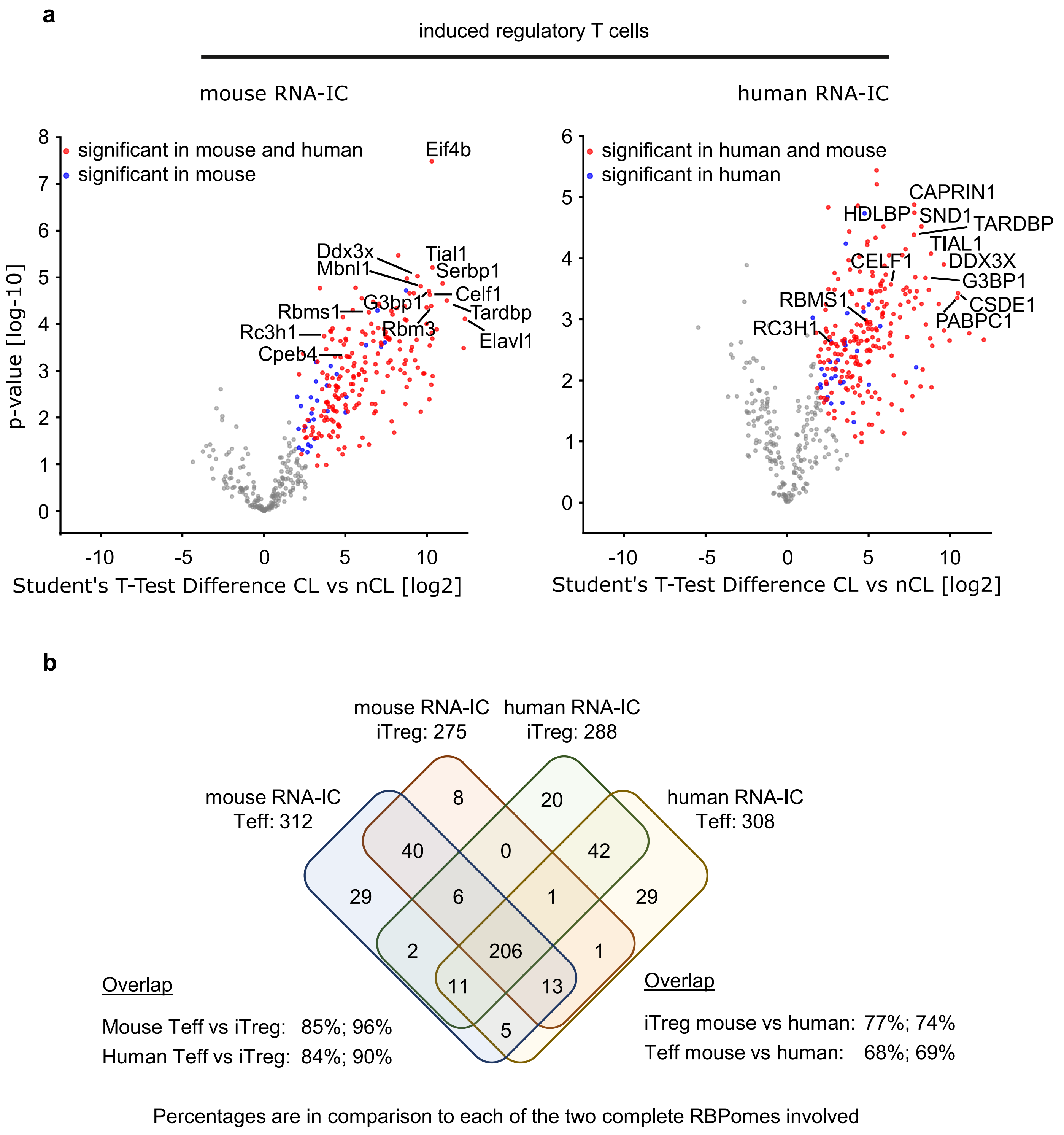

### Supplementary Figure 3

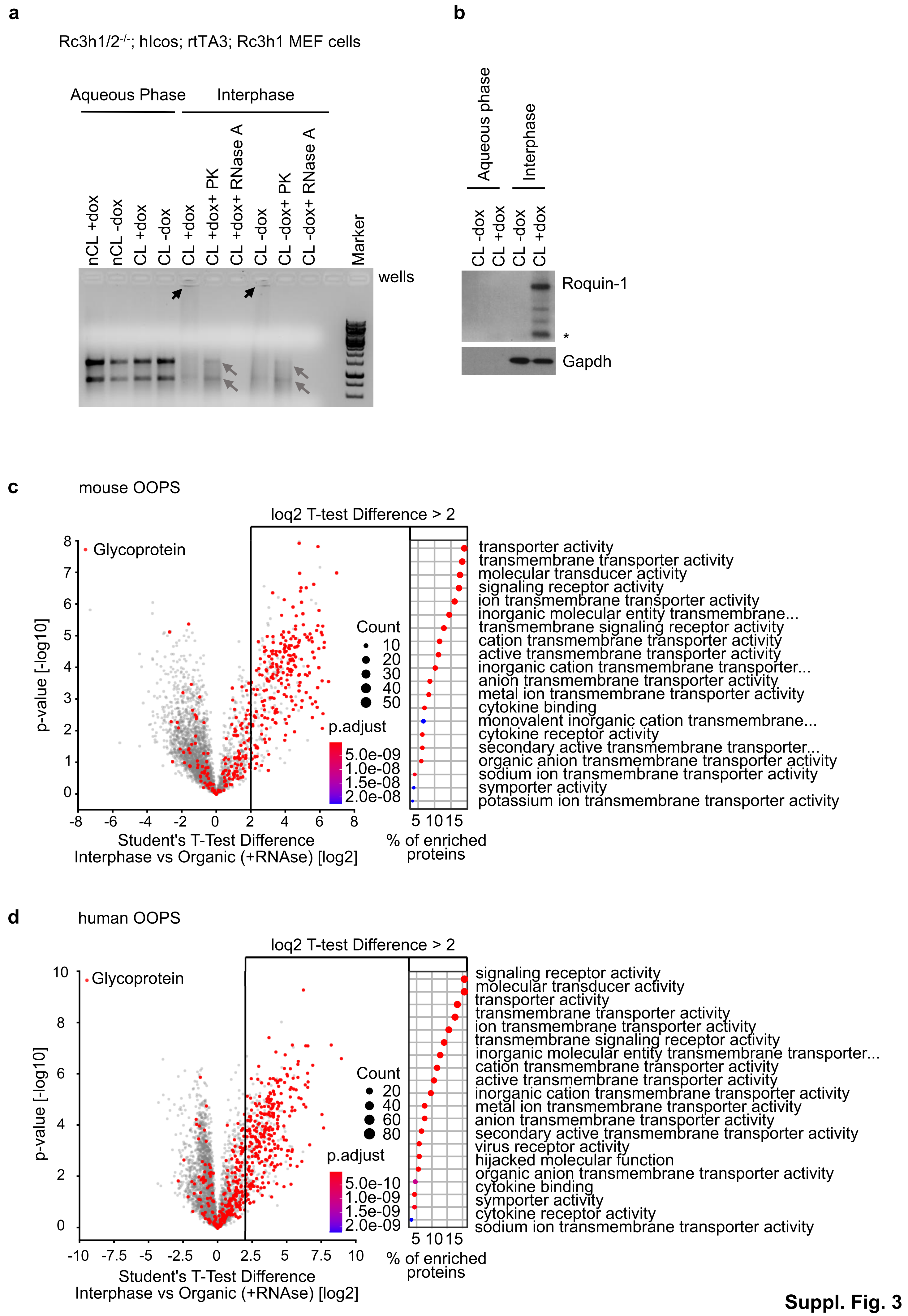

### Supplementary Figure 4

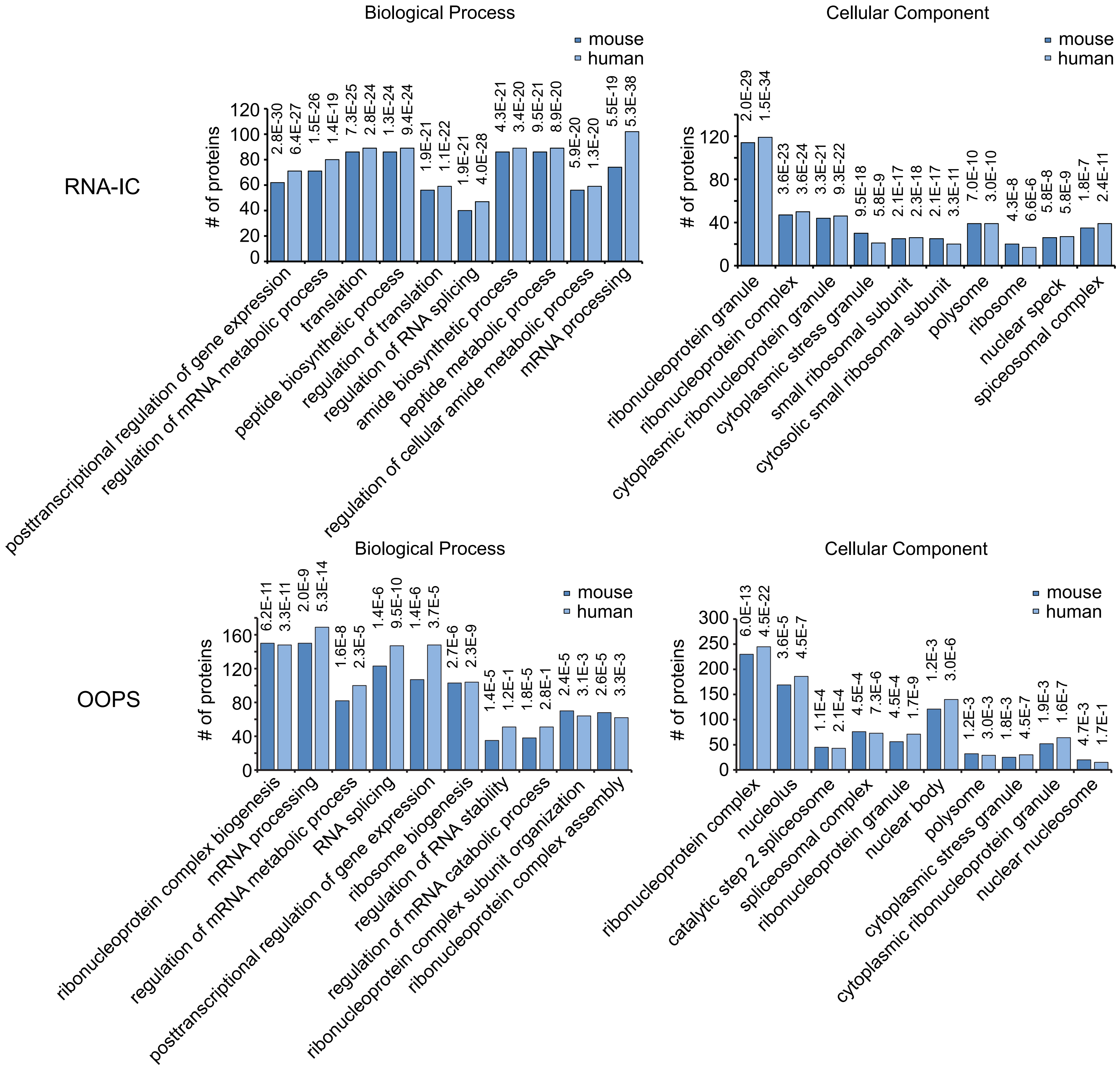

### Supplementary Figure 6

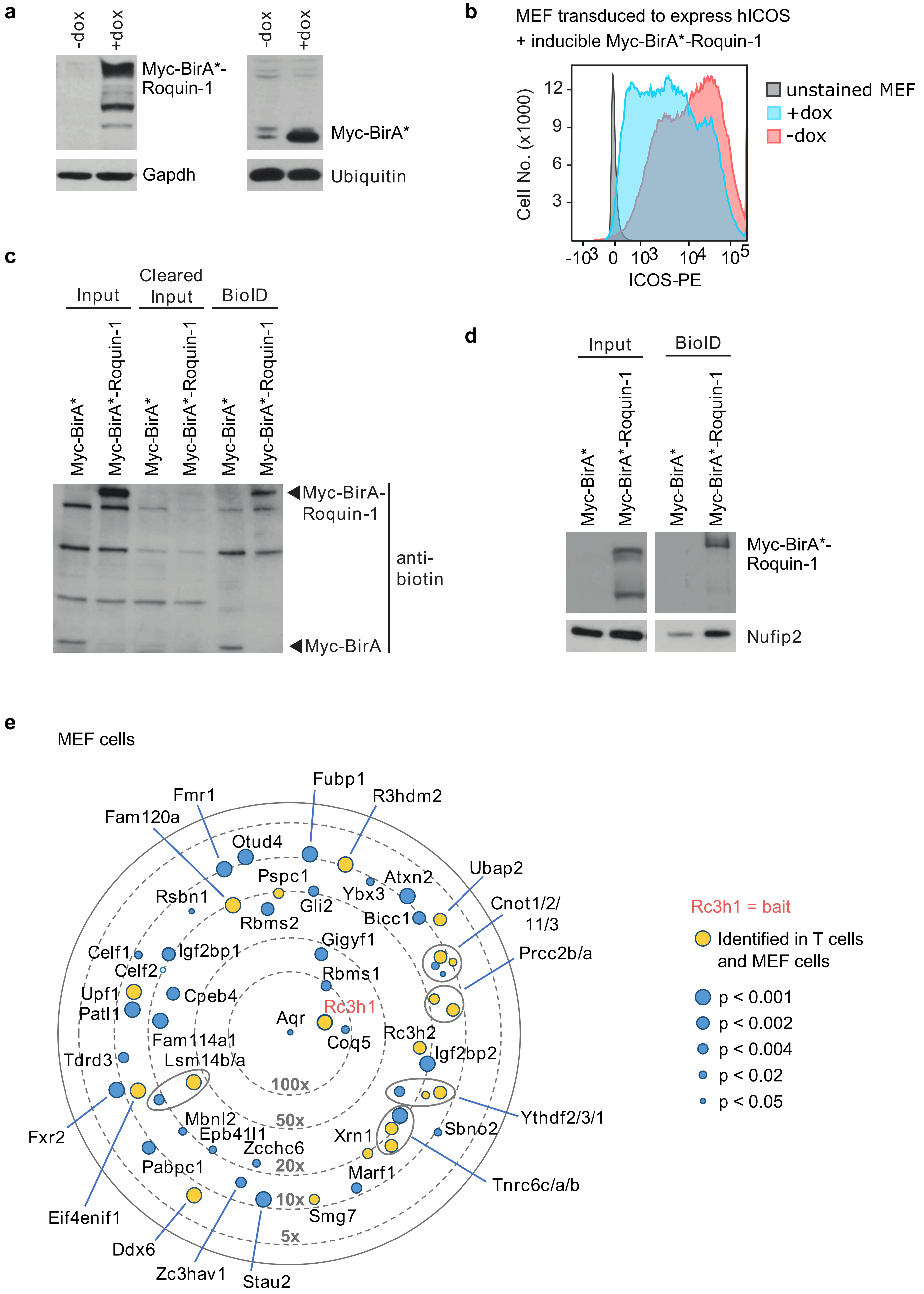

### Supplementary Figure 8

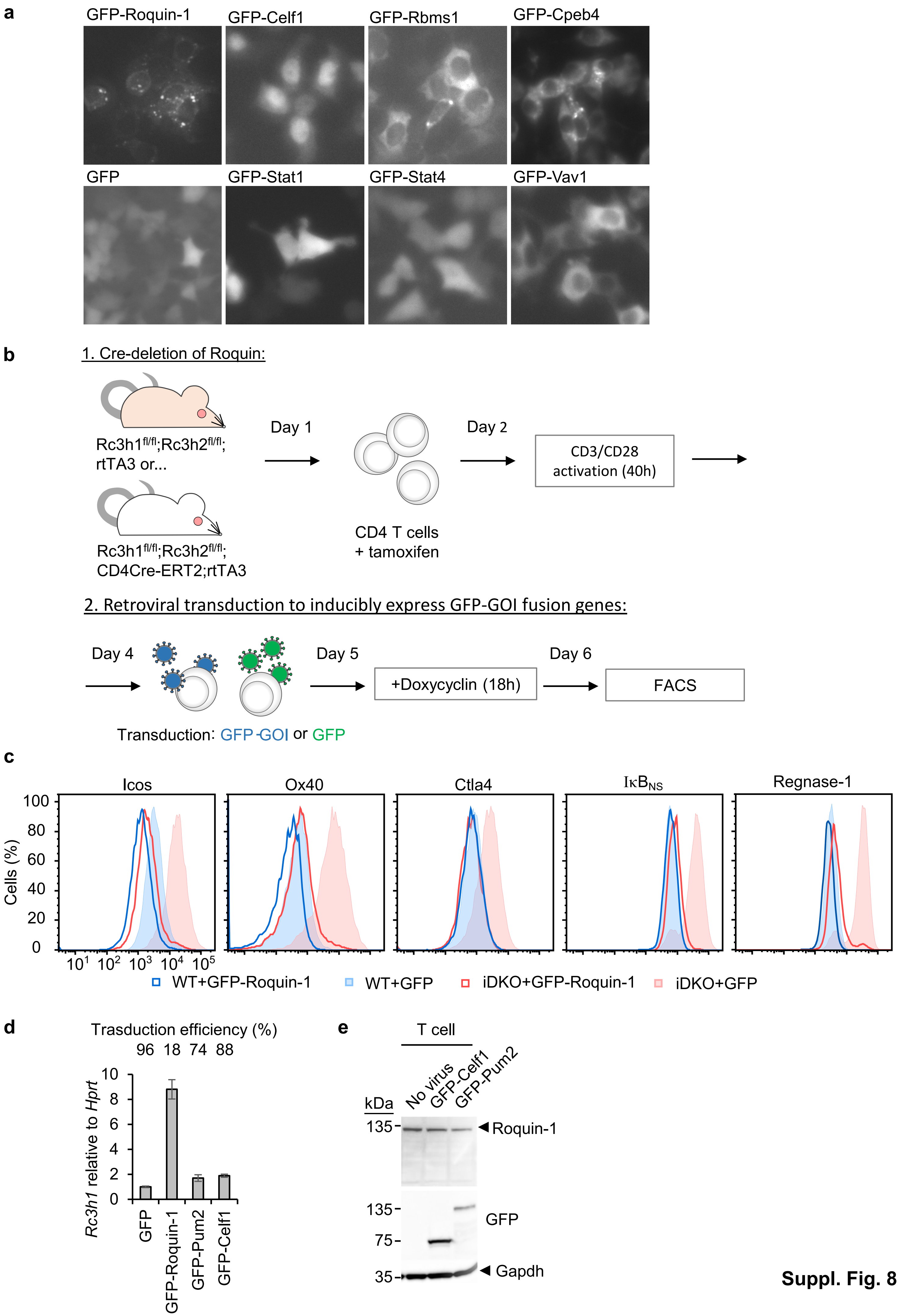
