## Supplementary Figure 5 for "Defining the RBPome of T helper cells to study higher order post-transcriptional gene regulation"

### zf-CCCH

| all proteins with annotated zf-CCCH domains | | | |
|---|---|---|---|
| Mus musculus | | Homo sapiens | |
| RNA-IC | OOPS | RNA-IC | OOPS |
| Cpsf4 | Cpsf4 | CPSF4 | CPSF4 |
| Cpsf4l | Cpsf4l | CPSF4L | CPSF4L |
| Dhx57 | Dhx57 | DHX57 | DHX57 |
| Helz | Helz | HELZ | HELZ |
| Leng9 | Leng9 | | |
| Mbnl1 | Mbnl1 | MBNL1 | MBNL1 |
| Mbnl2 | Mbnl2 | MBNL2 | MBNL2 |
| Mbnl3 | Mbnl3 | MBNL3 | MBNL3 |
| Mkrn1 | Mkrn1 | MKRN1 | MKRN1 |
| Mkrn2 | Mkrn2 | MKRN2 | MKRN2 |
| Mkrn3 | Mkrn3 | | |
| Parp12 | Parp12 | PARP12 | PARP12 |
| Ppp1r10 | Ppp1r10 | PPP1R10 | PPP1R10 |
| Prr3 | Prr3 | PRR3 | PRR3 |
| Rbm27 | Rbm27 | RBM27 | RBM27 |
| Rc3h1 | Rc3h1 | RC3H1 | RC3H1 |
| Rc3h2 | Rc3h2 | RC3H2 | RC3H2 |
| Rnf113a1 | Rnf113a1 | RNF113A | RNF113A |
| Rnf113a2 | Rnf113a2 | RNF113B | RNF113B |
| Toe1 | Toe1 | TOE1 | TOE1 |
| Trmt1 | Trmt1 | TRMT1 | TRMT1 |
| U2af1 | U2af1 | U2AF1 | U2AF1 |
| U2af1l4 | U2af1l4 | U2AF1L4 | U2AF1L4 |
| | | U2AF1L5 | U2AF1L5 |
| Unk | Unk | UNK | UNK |
| Unkl | Unkl | UNKL | UNKL |
| Zc3h10 | Zc3h10 | ZC3H10 | ZC3H10 |
| Zc3h13 | Zc3h13 | ZC3H13 | ZC3H13 |
| Zc3h15 | Zc3h15 | ZC3H15 | ZC3H15 |
| | | ZC3H18 | ZC3H18 |
| Zc3h3 | Zc3h3 | ZC3H3 | ZC3H3 |
| Zc3h4 | Zc3h4 | ZC3H4 | ZC3H4 |
| Zc3h6 | Zc3h6 | ZC3H6 | ZC3H6 |
| Zc3h7b | Zc3h7b | ZC3H7A | ZC3H7A |
| | | ZC3H7B | ZC3H7B |
| Zc3h8 | Zc3h8 | ZC3H8 | ZC3H8 |
| Zfp36 | Zfp36 | ZFP36 | ZFP36 |
| Zfp36l1 | Zfp36l1 | ZFP36L1 | ZFP36L1 |
| Zfp36l2 | Zfp36l2 | ZFP36L2 | ZFP36L2 |
| Zfp36l3 | Zfp36l3 | | |
| Zmat5 | Zmat5 | ZMAT5 | ZMAT5 |
| Zrsr1 | Zrsr1 | | |
| Zrsr2 | Zrsr2 | ZRSR2 | ZRSR2 |

## KH

| all proteins with annotated KH domains | | | |
|---|---|---|---|
| Mus musculus | | Homo sapiens | |
| RNA-IC | OOPS | RNA-IC | OOPS |
| 4921511C20 Rik | 4921511C20 Rik | AKAP1 | AKAP1 |
| Akap1 | Akap1 | ANKHD1 | ANKHD1 |
| Ankhd1 | Ankhd1 | ANKHD1-EIF4EBP3 | ANKHD1-EIF4EBP3 |
| Ankrd17 | Ankrd17 | ANKRD17 | ANKRD17 |
| Ascc1 | Ascc1 | ASCC1 | ASCC1 |
| Bicc1 | Bicc1 | BICC1 | BICC1 |
| Ddx43 | Ddx43 | DDX43 | DDX43 |
| | | DDX53 | DDX53 |
| Fmr1 | Fmr1 | FMR1 | FMR1 |
| Fubp1 | Fubp1 | FUBP1 | FUBP1 |
| Fubp3 | Fubp3 | FUBP3 | FUBP3 |
| Fxr1 | Fxr1 | FXR1 | FXR1 |
| Fxr2 | Fxr2 | FXR2 | FXR2 |
| Gm382 | Gm382 | | |
| Hdlbp | Hdlbp | HDLBP | HDLBP |
| Hnrnpk | Hnrnpk | HNRNPK | HNRNPK |
| Igf2bp1 | Igf2bp1 | IGF2BP1 | IGF2BP1 |
| Igf2bp2 | Igf2bp2 | IGF2BP2 | IGF2BP2 |
| Igf2bp3 | Igf2bp3 | IGF2BP3 | IGF2BP3 |
| Khdrbs1 | Khdrbs1 | KHDRBS1 | KHDRBS1 |
| Khdrbs2 | Khdrbs2 | KHDRBS2 | KHDRBS2 |
| Khdrbs3 | Khdrbs3 | KHDRBS3 | KHDRBS3 |
| Khsrp | Khsrp | KHSRP | KHSRP |
| Mex3a | Mex3a | MEX3A | MEX3A |
| Mex3b | Mex3b | MEX3B | MEX3B |
| Mex3c | Mex3c | MEX3C | MEX3C |
| Mex3d | Mex3d | MEX3D | MEX3D |
| Nova1 | Nova1 | NOVA1 | NOVA1 |
| Nova2 | Nova2 | NOVA2 | NOVA2 |
| Pcbp1 | Pcbp1 | PCBP1 | PCBP1 |
| Pcbp2 | Pcbp2 | PCBP2 | PCBP2 |
| Pcbp3 | Pcbp3 | PCBP3 | PCBP3 |
| Pcbp4 | Pcbp4 | PCBP4 | PCBP4 |
| Pnpt1 | Pnpt1 | PNPT1 | PNPT1 |
| Qk | Qk | QKI | QKI |
| Sf1 | Sf1 | | |
| Tdrkh | Tdrkh | TDRKH | TDRKH |

: canonical RBPs that were significantly enriched by the respective method

**Suppl. Fig. 5**
