## Supplementary Figure 7 for "Defining the RBPome of T helper cells to study higher order post-transcriptional gene regulation"

**a**

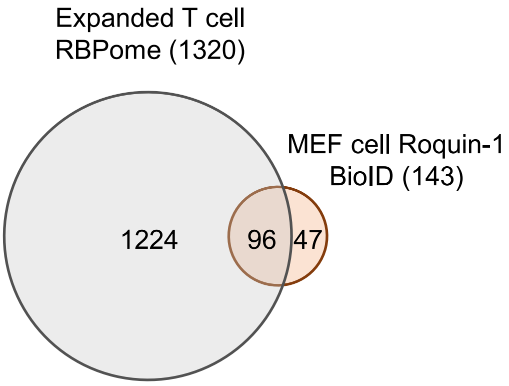


**b**

| | Roquin-1 MEF cell BioID overlap with RBPome | | | | |
|---|---|---|---|---|---|
| 1 | Acaca | FAM120A | Noc3l | R3hdm2 | Tdrd3 |
| 2 | Ankrd17 | Fmr1 | Nop16 | Rbms1 | Tnpo1 |
| 3 | Aqr | Fubp1 | Nop56 | Rbms2 | Tnrc6a |
| 4 | Atxn2 | Fxr1 | Nufip2 | Rc3h1 | Tnrc6b |
| 5 | Atxn2l | Fxr2 | Nup98 | Riok1 | Tnrc6c |
| 6 | Caprin1 | G3bp1 | Otud4 | Rpl23a | Ubap2 |
| 7 | Celf1 | Gigyf2 | Pabpc1 | Rpl26 | Ubap2l |
| 8 | Celf2 | Gnb2l1 | Patl1 | Rpl27a | Upf1 |
| 9 | Cenpe | Gnl3 | Pc | Rpl6 | Xrn1 |
| 10 | Cnot1 | Hist1h1c | Pds5b | Rpl8 | Ybx1 |
| 11 | Cnot11 | Igf2bp3 | Picalm | Rps14 | Ybx3 |
| 12 | Cnot2 | Kdm3b | Plec | Rps26 | Ythdf1 |
| 13 | Cnot3 | Kif1c | Prdx4 | Rsl1d1 | Ythdf2 |
| 14 | Cpeb4 | Larp4 | Prrc2a | Smg7 | Ythdf3 |
| 15 | Csde1 | Larp4b | Prrc2b | Snw1 | Zc3hav1 |
| 16 | Ddx18 | Lsm14a | Prrc2c | Stat4 | Zcchc6 |
| 17 | Ddx27 | Lsm14b | Pspc1 | Stau1 | |
| 18 | Ddx6 | Lyar | Ptbp1 | Strap | |
| 19 | Dhx9 | Marf1 | Pum1 | Syncrip | |
| 20 | Eif4enif1 | Mbnl2 | Pum2 | Tardbp | |

☐ cloned for induced expression; detected in T and MEF cell BioIDs

☐ cloned for induced expression; detected in MEF cell BioID only

**Suppl. Fig. 7**
